# DISCOVERY AND VALIDATION OF GENES DRIVING DRUG-INTAKE AND RELATED BEHAVIORAL TRAITS IN MICE

**DOI:** 10.1101/2023.07.09.548280

**Authors:** Tyler A. Roy, Jason A. Bubier, Price E. Dickson, Troy D. Wilcox, Juliet Ndukum, James W. Clark, Stacey J. Sukoff Rizzo, John C. Crabbe, James M. Denegre, Karen L. Svenson, Robert E. Braun, Vivek Kumar, Stephen A. Murray, Jacqueline K. White, Vivek M. Philip, Elissa J. Chesler

**Affiliations:** The Jackson Laboratory, Bar Harbor, ME; Joan C Edwards School of Medicine, Marshall University Huntington, WV; University of Pittsburgh School of Medicine, Pittsburgh, PA; Oregon Health & Science University and VA Portland Health Care System, Portland, OR

## Abstract

Substance use disorders (SUDs) are heritable disorders characterized by compulsive drug use, but the biological mechanisms driving addiction remain largely unknown. Genetic correlations reveal that predisposing drug-naïve phenotypes, including anxiety, depression, novelty preference, and sensation seeking, are predictive of drug-use phenotypes, implicating shared genetic mechanisms. Because of this relationship, high-throughput behavioral screening of predictive phenotypes in knockout (KO) mice allows efficient discovery of genes likely to be involved in drug use. We used this strategy in two rounds of screening in which we identified 33 drug-use candidate genes and ultimately validated the perturbation of 22 of these genes as causal drivers of substance intake. In our initial round of screening, we employed the two-bottle-choice paradigms to assess alcohol, methamphetamine, and nicotine intake. We identified 19 KO strains that were extreme responders on at least one predictive phenotype. Thirteen of the 19 gene deletions (68%) significantly affected alcohol use three methamphetamine use, and two both. In the second round of screening, we employed a multivariate approach to identify outliers and performed validation using methamphetamine two-bottle choice and ethanol drinking-in-the-dark protocols. We identified 15 KO strains that were extreme responders across the predisposing drug-naïve phenotypes. Eight of the 15 gene deletions (53%) significantly affected intake or preference for three alcohol, eight methamphetamine or three both (3). We observed multiple relations between predisposing behaviors and drug intake, revealing many distinct biobehavioral processes underlying these relationships. The set of mouse models identified in this study can be used to characterize these addiction-related processes further.

## 1 INTRODUCTION

Substance use disorders (SUDs) are highly heritable and widely prevalent brain diseases^1^ that manifest themselves both behaviorally and physiologically ^2,3^. Currently, over 20 million people ages twelve and up are suffering from a SUD in the United States^4^, and drug and alcohol use costs Americans more than $700 billion and contributes to 570,000 deaths per year^2,5^.

Despite extensive efforts to identify and characterize mechanisms driving substance use, few pharmacotherapeutic treatments exist.^3^ This may be due, at least partly, to a historical emphasis on the deep characterization of a few well-known biological mechanisms influencing substance use rather than discovering novel and perhaps unexpected genetic mechanisms influencing substance use. Due to the conservation of many aspects of the addiction-related reward circuitry across species^6,7^, it is possible to leverage the exquisite resources of mouse genetics to discover new biological mechanisms of addiction risk behaviors^6,7^.

Genetic and genomic screens have been previously employed in mutant mouse strains to identify novel addiction risk mutations. A major challenge in these studies is that they require a separate drug-exposed cohort of mice to avoid the effects of drug exposure on subsequent physiology and behaviors^8^. Further, procedures designed to screen candidates for their role in addiction-related phenotypes (i.e., initiation, escalation, extinction, reinstatement) require the application of expensive and complex experimental paradigms to characterize drug consumption and drug effects in mice. In more recent high-throughput, discovery-based approaches conducted by the International Mouse Phenotyping Consortium (IMPC), large-scale screens which employ a single unified test battery were found to efficiently characterize behavioral and physiological phenotypes of single-gene C57BL/6NJ KO strains^9^. These results suggests that large-scale behavioral screens that include predictive phenotypes as part of the test batteries might efficiently identify novel addiction-related genes and pathways.

Many risk factors for and consequences of addiction and SUDs are associated with other predisposing drug-naïve phenotypes, personality traits, and co-occurring psychological conditions in humans, including anxiety, depression, impulsivity, and novelty-seeking^10-13^. Using mouse behavioral tests, it is possible to precisely model many aspects of these predisposing or co-occurring traits^13-15^. Previous rodent studies have shown that predisposing drug-naïve phenotypes, which can be assayed using approach-avoidance tasks, ‘behavioral despair’ assays, and novelty-seeking tasks, can be used to predict future drug-related behavioral phenotypes, such as conditioned place preference, sensitization, and self-administration^16-19^. Additional studies using inbred mouse populations have revealed shared genetic mechanisms driving predisposing drug-naïve phenotypes and drug-related behavioral phenotypes across distinct drug classes^20,21^. However, despite these efforts, many genes underlying the shared genetic variation among drugs, alcohol, and predisposing drug-naïve phenotypes remain unknown. In the present study, we exploited these relationships to identify novel addiction risk genes by screening single-gene KO strains and evaluated whether these strains exhibited altered consumption or preference for alcohol and methamphetamine.

## 2 MATERIALS AND METHODS

### 2.1 Animal Care and Husbandry

Mice in all experiments were maintained in a climate-controlled room under a standard 12:12 light-dark cycle (lights on at 0600 hours and off at 1800 hours). They were provided free access to food (NIH315K52 chow, Lab Diet 6%/PM Nutrition, St. Louis, MO, USA) and acidified water with vitamin K supplementation unless indicated otherwise. All husbandry, procedures, and protocols were approved by The Jackson Laboratory (JAX) Animal Care and Use Committee and were conducted in compliance with the National Institutes of Health Guidelines for Care and Use of Laboratory Animals. All details for housing and testing conditions can be found at https://www.mousephenotype.org/impress/PipelineInfo?id=12.

We followed JAX’s rigorous genetic quality control and mutant gene genotyping programs so that the genetic background and integrity of the mutation were maintained. In addition to the quality control JAX employs to maintain the integrity of the background strains, these quality control measures were also employed to maintain the integrity of the genotypes of strains with identified molecular mutations. For example, all KO strains used in this project were created using C57BL/6NJ (RRID: IMSR_JAX:005304) embryonic stem cells such that no flanking DNA differs from controls and mutants. Similarly, all endonuclease-modified strains used have no flanking DNA, which differs from control strains. In addition, we received all strains for our screens directly from JAX production colonies at wean, ensuring that all strains tested met requirements for rigorous genetic quality control of background and mutations.

### 2.2 Overview of Behavioral Phenotyping Procedures

The Knockout Mouse Project (KOMP) phenotyping center (KOMP2, RRID: SCR_017528) at JAX was established in 2011 to generate and phenotype 833 single-gene knockout (KO) mouse strains. The KOMP2 pipeline includes measures of physiological, behavioral, and biochemical characteristics and the implementation of a standardized battery of analyses to characterize the effects of gene KOs. We performed analyses on measured traits using the R/PhenStat Bioconductor package (v 1.0.0)^22^. PhenStat (RRID: SCR_021317) is built on a linear mixed-effects model where the date of the test is considered the random effect with sex, genotype, and the interaction of sex and genotype information as fixed effects terms. Missing values were ignored.

Using data collected from the KOMP2 pipeline, we undertook two rounds of screening for KO strains for phenodeviance on predisposing drug-naïve phenotypes and subsequent drug-use evaluation, the first in 2014 and the second in 2017. Within each cohort, mice were underwent the full battery of tests of biological and behavioral endpoints, including assays such as glucose tolerance, open field and light-dark ^9^. Tests were arranged in a fixed order by perceived stressfulness to minimize potential carry-over effects (Fig. S1). In addition, all runs within the phenotyping pipeline were conducted in a sex-specific manner, e.g., each run consisted exclusively of males or females. Protocols for all tests can be found at https://www.mousephenotype.org/impress/PipelineInfo?id=7.

### 2.3 Cohort #1: Relationship of Predisposing Drug-Naive Phenotypes to Drug-Intake Phenotypes: Phenodeviance, Two-Bottle Choice and Principal Component Analysis

Within the KOMP2 resource, 221 KO strains had undergone behavioral phenotyping at the time of the first screen in 2014. These strains were matched with C57BL/6NJ controls and tested on a broad behavioral phenotyping pipeline that included as part of the KOMP2 project (Fig. S1) five behavioral assays (tail suspension, acoustic startle, open field, light/dark, and hole board) that define ten predisposing drug-naïve phenotypes (Table S1) previously shown to predict drug-related behaviors in mice^14,15,23-25^. Protocols for the tail suspension tests^25^, acoustic startle^26^, open field^19^, light/dark^14^, and hole board^27^ match the SOPs used at the time of testing (2014).

#### 2.3.1 Detecting Predisposing Drug-Naïve Phenodeviance

We rankZ transformed the data from 221 KO strains and analyzed it by the linear mixed model within PhenStat (v 1.0.0)^22,28^. We found 143 significantly phenodeviant strains from C57BL/6NJ controls (p<0.05) on at least one of the ten chosen predisposing drug-naïve phenotypes and identified them as extreme strains. We prioritized strains with multiple significant predisposing phenotypes for further testing for drug-use phenotypes; however, testing was restricted to strains available at the time of the study. Of the 143 strains that were phenodeviant on at least one of the predisposing traits, 19 were selected for further testing because they existed as established live colonies (rather than frozen embryos) and were thus available for test cohort production. The 19 strains included: C57BL/6NJ-*Btg2*^tm1b(*KOMP*)Mbp^/2J, B6N(Cg)-*C1qa^tm1b(EUCOMM)Wtsi^*/3J, B6N(Cg)-*C9^tm1.1(KOMP)Vlcg^*/J, B6N(Cg)-*Cfb^tm1.1(KOMP)Wtsi^*/J, B6N(Cg)-*Cp^tm1b(KOMP)Wtsi^*/J, B6N(Cg)-*Dnajb3^tm1.1(KOMP)Vlcg^*/J, C57BL/6NJ-*Dnase1l2^em1(IMPC)J^*/J, B6N(Cg)-*Epb41l4a^tm1b(KOMP)Mbp^*/2J, B6N(Cg)-*Far2^tm2b(KOMP)Wtsi^*/2J, B6N(Cg)-*Gipc3^tm1b(KOMP)Wtsi^*/J, B6N(Cg)-*Hdac10^tm1.1(KOMP)Mbp^*/J, B6N(Cg)-*Hspb2^tm1.1(KOMP)Vlcg^*/J, B6N(Cg)-*Htr1a^tm1.1(KOMP)Vlcg^*/J B6N(Cg)-*Il12rb2^tm1.1(KOMP)Vlcg^*/J, B6N(Cg)-*Lpar6^tm1.1(KOMP)Vlcg^*/J, B6N(Cg)-*Parp8^tm1.1(KOMP)Wtsi^*/J, B6N(Cg)-*Pitx3^tm1.1(KOMP)Vlcg^*/J, B6N(Cg)-*Pnmt^tm1.1(KOMP)Vlcg^*/J, B6N(Cg)-*Rilpl2^tm1b(KOMP)Wtsi^*/J. All strains were homozygous for their gene deletions. All mutants were tested relative to sex and age-matched control C57BL/6NJ mice.

#### 2.3.2 Two-Bottle Choice Assay to Evaluate Drug-Related Phenotypes

Mice from the 19 KO strains selected for the two-bottle choice (2BC) protocol were obtained from the JAX Repository and transferred to the JAX housing and phenotyping facility. Mice were group-housed, with no more than five of the same sex, in duplex polycarbonate cages before testing. Using a 2BC assay, we then determined drug-use phenotypes by defining substance use for ethanol (EtOH), nicotine, or methamphetamine (MA) in mice from each of these strains (see Table S2 for sample sizes).

At a minimum of one day before testing, we rehoused the mice individually in duplex polycarbonate cages with a single Shepherd Shack® and Nestlet® for the duration of testing. We kept the single housing time minimal to reduce the effects of social isolation^29^. The 2BC protocol was adapted from one previously published^30^ to test three different drugs at varying concentrations: EtOH (3%, 6%, 12%, and 15%), nicotine (10 mg/L, 20 mg/L, 40 mg/L, and 80 mg/L), and MA (10 mg/L, 20 mg/L, 40 mg/L, and 80 mg/L). All three drugs were diluted in sterilized acidified (pH 2.5-3) water. The nicotine solution also contained 20 g/L saccharine sodium salt hydrate to mitigate the bitter taste. For each drug, mice were exposed to both a tube of water and a tube of the drug at the indicated concentration. Each dose was tested for two days before switching to the next dose which was 2x the prior dose. Individual mice were exposed to only one drug for testing. From these data, we obtained measures of drug preference and drug consumption. Drug preference was defined as the volume of drug consumed/ total fluid volume consumed (drug + water), whereas drug consumption is defined as the amount of drug consumed (mL of drug consumed × mg/mL drug)/ kg body weight. We also analyzed water intake and per total fluid intake as these measures facilitate the interpretation of drug preference and consumption outcomes. Water intake is the total volume of pure water ingested over the specified time frame, whereas total fluid intake is the total volume of pure water plus drug solution consumed over the specified time frame. Due to strain availability at the time of testing, we tested 16 of the 19 strains with all three drugs; EtOH data is missing for one strain [*C9*], while nicotine and MA data are missing for two strains [*Lpar6* and *Pnmt*].

We then applied a repeated-measures ANOVA to each of the 2BC phenotypes to assess the strain × sex × dose effects and evaluate strain and strain × sex effects. After fitting each model, we obtained the least-squares mean difference between each KO relative to the C57BL/6NJ controls. We used a threshold of false discovery rate (FDR) < 0.05 to test for significance of effects in the model.

#### 2.3.3 Principal Component Analysis to Define Relationships Among Phenotypes

To investigate the underlying shared correlation structure across drug naïve and drug-use behaviors, we conducted a principal component analysis (PCA)^31^. All genotype difference estimates were subjected to a Van der Weerden (RankZ) transformation and PCA in R (V 3.4.4) using factoextra _1.0.5, fviz_pca_biplot (RRID:SCR_016692)^32^. We extracted the first two principal components (PCs) to assess the relationships among the ten predisposing drug-naïve phenotypes, the six drug-use 2BC traits (consumption, preference x three drugs), and the six liquid consumption 2BC traits (water, total liquid intake x three drugs). We then performed a PCA biplot clustering using the effect sizes across predisposing drug-naïve phenotypes and 2BC traits. Because PCA can only be conducted on complete data sets and because *C9, Lpar6,* and *Pnmt* data were incomplete, we only analyzed 16 of the 19 KO strains by PCA.

### 2.4 Cohort #2: Relationship of Predisposing Drug-Naive Phenotypes to Drug-Use Phenotypes: Multivariate Outlier Detection, Two-Bottle Choice, and Drinking in the Dark Assay

We took advantage of the ongoing KOMP program for our second test cohort. At the time we identified our second validation cohort, a total of 401 KO strains had undergone behavioral phenotyping. These 401 strains included all 221 of the strains that were included in the first cohort selection. Mice from these strains were matched with temporally local C57BL/6NJ controls and tested on four of the five behavioral assays (acoustic startle, open field, light/dark, and hole board) that define eight of the ten predisposing drug-naïve phenotypes that were used in Cohort #1 (Fig. S1). The tail suspension assay, including measures of time immobile and latency to immobility phenotypes, was dropped from KOMP2 testing and was excluded from the strain identification as this missing data would have greatly reduced the number of strains with sufficient data for the Cohort #2 analysis.

#### 2.4.1 Mahalanobis Distance to Identify Predisposing Drug-Naïve Phenotypes

In this cohort, we used the Mahalanobis distance to identify which of the 401 KO strains were phenodeviant across the eight predisposing phenotypes. We chose this approach because our initial cohort revealed non-uniform, multidimensional relations underlying the drug use and their predisposing drug-naïve phenotypes. Mahalanobis identifies multivariate outliers strains by calculating the distance from the centroid, representative of control strain values, in a multidimensional space^33^. The centroid is defined as the intersection of the mean of the variables being assessed. The Mahalanobis distance follows a *χ*^2^ distribution, which is used to evaluate statistical significance^33^. This was used to create one score representing the combined phenodeviance across all eight predisposing phenotypes. We used the R/Phenstat Bioconductor package (v 1.0.0)^22^ for modeling the association between trait and genotype. We then performed a rankZ transformation and input the transformed genotype effect estimates to Mahalanobis distance calculations. Phenodevience was defined as a combined Mahalanobis score that was significantly extreme based on *χ*^2^; using this criterion, we identified 123 of the 401 strains as significantly phenodeviant from the C57BL/6NJ controls.

Of these 123 phenodeviant strains identified, our goal was to rederive and test further the most extreme 25 strains as defined by their highest scores. Due to the availability of sperm, the success of *in vitro* fertilization, and the ability to produce viable cohorts from each strain, we tested 13 of these most extreme phenodeviant strains. We also included *Tmod2* and *Rap2b* KO strains, which were phenodeviant as determined by the Mahalanobis distance calculations but not in the top 25 most phenodeviant, due to expert recommendation. The 15 strains selected for the second cohort include: B6N(Cg)-Elof1^tm1.1(KOMP)Vlcg/J(+/-)^, C57BL/6NJ-Stk36^em1(IMPC)J^/J^(-/-)^, C57BL/6NJ-Myh10^em1(IMPC)J^/J^(+/-)^, B6N(Cg)-Dnmt3a^tm1b(KOMP)Wtsi^/J^(+/-)^, B6N(Cg)-Cp^tm1b(KOMP)Wtsi^/J^(-/-)^, B6N(Cg)-Zbtb4^tm1.1(KOMP)Vlcg^/J^(-/-)^, B6N(Cg)-Dnaja4^tm1b(KOMP)Wtsi^/J^(-/-)^, B6N(Cg)-Irf8^tm1b(KOMP)Wtsi^/J^(-/-)^, B6N(Cg)-Htr7^tm1b(KOMP)Wtsi^/J^(-/-)^, B6N(Cg)-Gpr142^tm1.1(KOMP)Vlcg^/J^(-/-)^, B6N(Cg)-C3^tm1.1(KOMP)Vlcg^/J^(-/-)^, B6N(Cg)-Stx19^tm1.1(KOMP)Vlcg^/2J^(-/-)^, B6N(Cg)-Lrrc15^tm1b(KOMP)Wtsi^/J^(-/-)^, B6N(Cg)-Rap2b^tm1.1(KOMP)Vlcg^/J^(+/-)^, and B6N(Cg)-Tmod2^tm1b(KOMP)Wtsi^/2J^(-/-)^. These strains were obtained from the JAX Breeding and Rederivation Services and transferred to the JAX housing and phenotyping facility, where they were bred to testable cohort sizes through pair and trio mating. Viable homozygous null strains were bred using -/- × -/- breeding pairs or trios (1M, 2F), whereas lethal homozygous null strains were bred using +/- × +/- breeding pairs or trios.

#### 2.4.2 Methamphetamine Two-Bottle Choice

We tested mice for drug-use phenotypes in Cohort #2 using the MA 2BC since we wanted to continue rapidly screening mice for MA-use phenotypes. We tested mice for MA use phenotypes using the same protocol described for Cohort #1, including the same four doses of MA (10 mg/L, 20 mg/L, 40 mg/L, and 80 mg/L). The number of mice tested for each KO strain is indicated in Table S3. We did not test for nicotine oral self-administration because we failed to find any effects using our nicotine protocol in Cohort #1.

#### 2.4.3 Ethanol Drinking in the Dark Protocol

We used the EtOH drinking-in-the-dark (DID) assay to screen mice for the more translationally relevant binge drinking phenotype. Although preference drinking (EtOH 2BC) is a widely used and valid partial model for alcohol use,^34,35^ the DID protocol has improved relevance as a model of binge drinking.^36-38^ It has been well documented that there is significant comorbidity of alcohol and MA use^30^. However, newer evidence suggests that binge drinking has a significantly higher comorbidity rate and is a better predictor for MA use than moderate drinking.^39,40^ Using these two protocols, we screened our extreme single-gene KO strains for EtOH binge drinking effects and strong multi-drug effects that have translational relevance to human patterns of MA and alcohol use.

We tested adult (8-24 week old) mice using the previously published EtOH DID protocol^36-38^. This protocol has been refined to induce mice to drink to levels of intoxication (∼100 mg/dL)^41,42^. The number of mice tested for each KO strain is indicated in Table S4. The EtOH DID protocol is a four-day, limited-access protocol in which EtOH is available during a time in the circadian cycle when mice are behaviorally active (nocturnal) to induce binge drinking behaviors that lead to intoxication on the final day. We applied a repeated-measures ANOVA to EtOH consumption across the four days of the DID protocol to assess the strain × sex effects, in addition to assessing strain effects. Following the model fit, we obtained the least squares mean difference between each KO relative to the C57BL/6NJ control. We used a threshold of FDR < 0.05 to determine the significance of terms in the model. Of particular interest were strain, and strain × sex effects.

#### 2.4.4 Blood Collection and Blood Ethanol Concentration Analysis

We collected a minimum of 50 µL of blood into a microtainer (VWR, cat.# VT365956) immediately following ethanol removal on the final day of the DID protocol. We centrifuged blood samples at 13,300 RPM for 11 minutes and pipetted serum into separate Eppendorf tubes on dry ice within one hour of collection. We transferred the serum samples to a -80°C freezer for storage within three hours of collection. We determined blood ethanol concentration (BEC) using a Beckman DXC biochemical analyzer (RRID:SCR_019633). We analyzed BEC using a two-way ANOVA and evaluated significant differences between each KO relative to the C57BL/6NJ control using Tukey’s Honestly Significant Difference test.

### 2.5 Functional Analysis of Candidate Genes

To evaluate whether and how the genes identified in our knockout analysis might be involved in addiction-related traits, we searched for genetic and genomic evidence to identify plausible biological mechanisms in which the genes could have affected drug-use phenotypes. To accomplish this, we performed a functional analysis of all the statistically significant genes using GeneWeaver (RRID:SCR_003009)^43^. We also conducted a systematic search of the genes which altered 2BC phenotypes to determine whether they were represented in previous curated genomic data sets from studies of humans, mice, and rats. Among the data resources used in the analysis were the following: (a) Medical Subject Headings (MeSH) (RRID:SCR_004750)^44^ related to drugs or addiction, (b) Gene Ontology (GO) (RRID:SCR_006447)^45,46^ terms related to drugs or addiction, (c) Quantitative Trait Loci (QTL)^47^ gene sets related to drugs or addiction, (d) Kyoto Encyclopedia of Genes and Genomes (KEGG) (RRID:SCR_001120)^48-50^ pathways related to addiction and alcoholism, (e) Neuroinformatics Framework Drug-Related Genes (DRG) (RRID:SCR_003330)^51^, and (f) Genome-Wide Association Studies (GWAS)^52^ of alcohol and substance use related traits. In addition, we performed a Gene Set Enrichment Analysis (GSEA, RRID:SCR_003199)^53^.

## 3 Results

### 3.1 Detecting Predisposing Drug-Naïve Phenodeviance in Cohort #1

Of the 221 KO strains tested, 143 (64.7%) were phenodeviant defined by at least one of the ten predisposing drug-naïve phenotypes being significantly different (p<0.05) from C57BL/6NJ controls. Of these 143 phenodeviant strains, we further analyzed 19 for drug-intake phenotypes (the remaining 124 strains were no longer available for testing as they had been cryopreserved and the strains no longer maintained). The rankZ results of the 10-predisposing drug-naïve phenotypes obtained from the battery of five behavioral tests for these 19 strains are shown in Figure 1. Thus, while the KO strains chosen to screen for drug-use phenotypes were limited due to availability, the 19 strains tested were nonetheless highly representative of the range of phenodeviance observed within the 143 phenodeviant stains for each predisposing drug-naïve trait indicating that there was not any inherent bias in the 19 strains tested for drug-use phenotypes.

**Figure 1:**
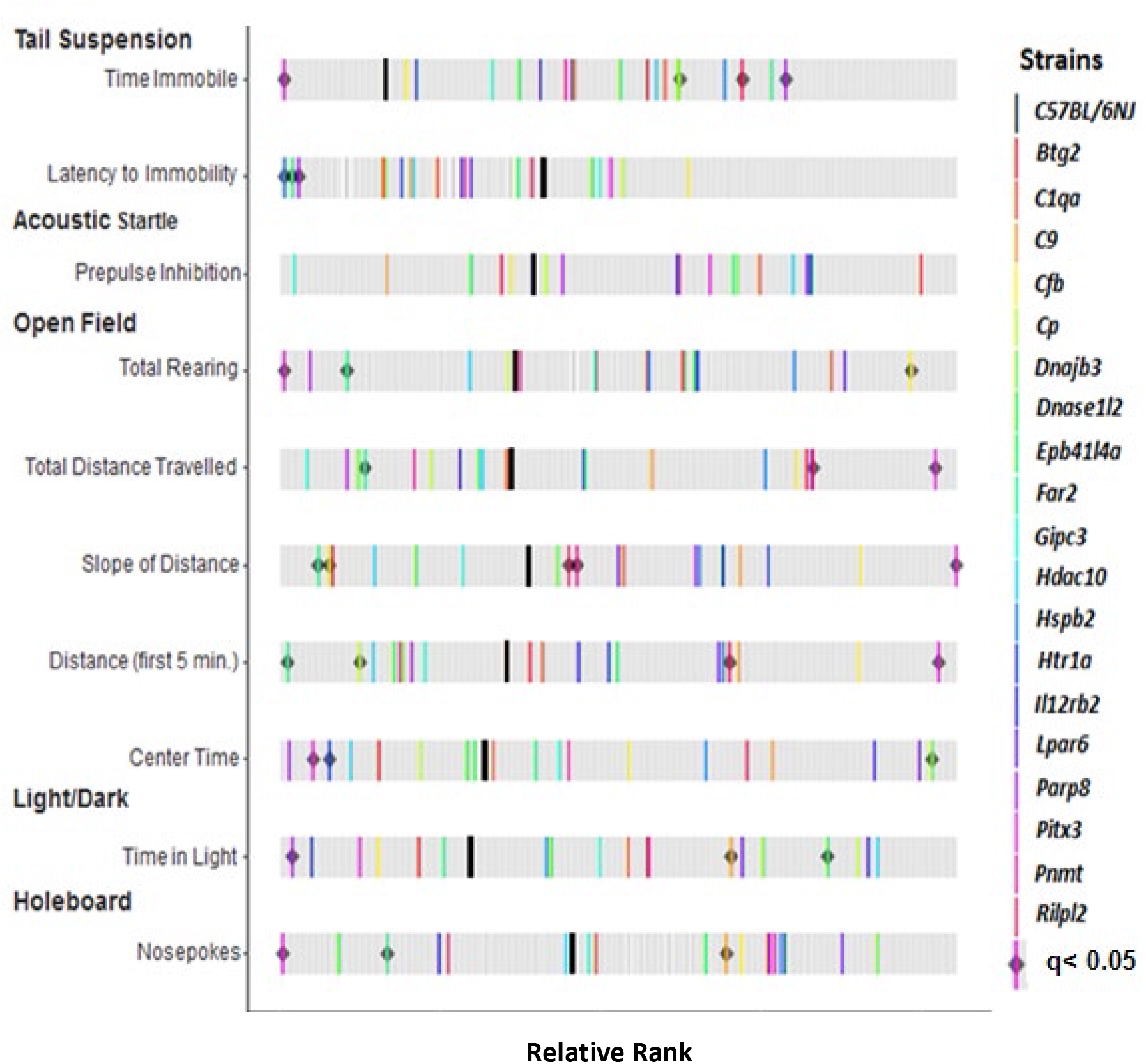
Phenotypic variation in predisposing drug-naïve phenotypes from the 19 phenodeviant strains identified in Cohort #1. Of the 221 strains tested in 2014, 143 strains were phenodeviant in at least one predisposing, drug-naïve phenotype, and 19 were established colonies and tested further for drug-use phenotypes using the two-bottle choice assay. The Rank Z graph displays where the 19 strains fall in the range of the 221 strains measured for each of the 10 predisposing, drug-naïve phenotypes. Thick black bars represent C57BL6/NJ controls. Colored bars represent strains identified as phenodeviant and predictive of addiction risk phenotypes. Gray bars represent the ranked genotype effects for each measure calculated across all 201 KO strains tested in Cohort #1. Black diamonds indicate KO strains from initial screening which remained significantly phenodeviant when accounting for multiple testing corrections (q < 0.05).

### 3.2 Two-Bottle Choice Assay to Determine Drug-Use Phenotypes

We tested these 19 single-gene KO strains using the 2BC test for drug consumption and preference for EtOH, nicotine, and MA for a total of six drug-use phenotypes. Based on significant effects by strain or strain × dose, we found that 15 of the 19 strains showed at least one significant drug-use effect; only *Epb41l4a, Pitx3, Gipc3*, and *C9* showed no significant differences from controls in any of the six drug-use phenotypes tested (Fig. 2A). In contrast, *Il12rb* and *Far2* each showed significant differences from controls for three of the six phenotypes, the most of all 19 strains tested.

**Figure 2.**
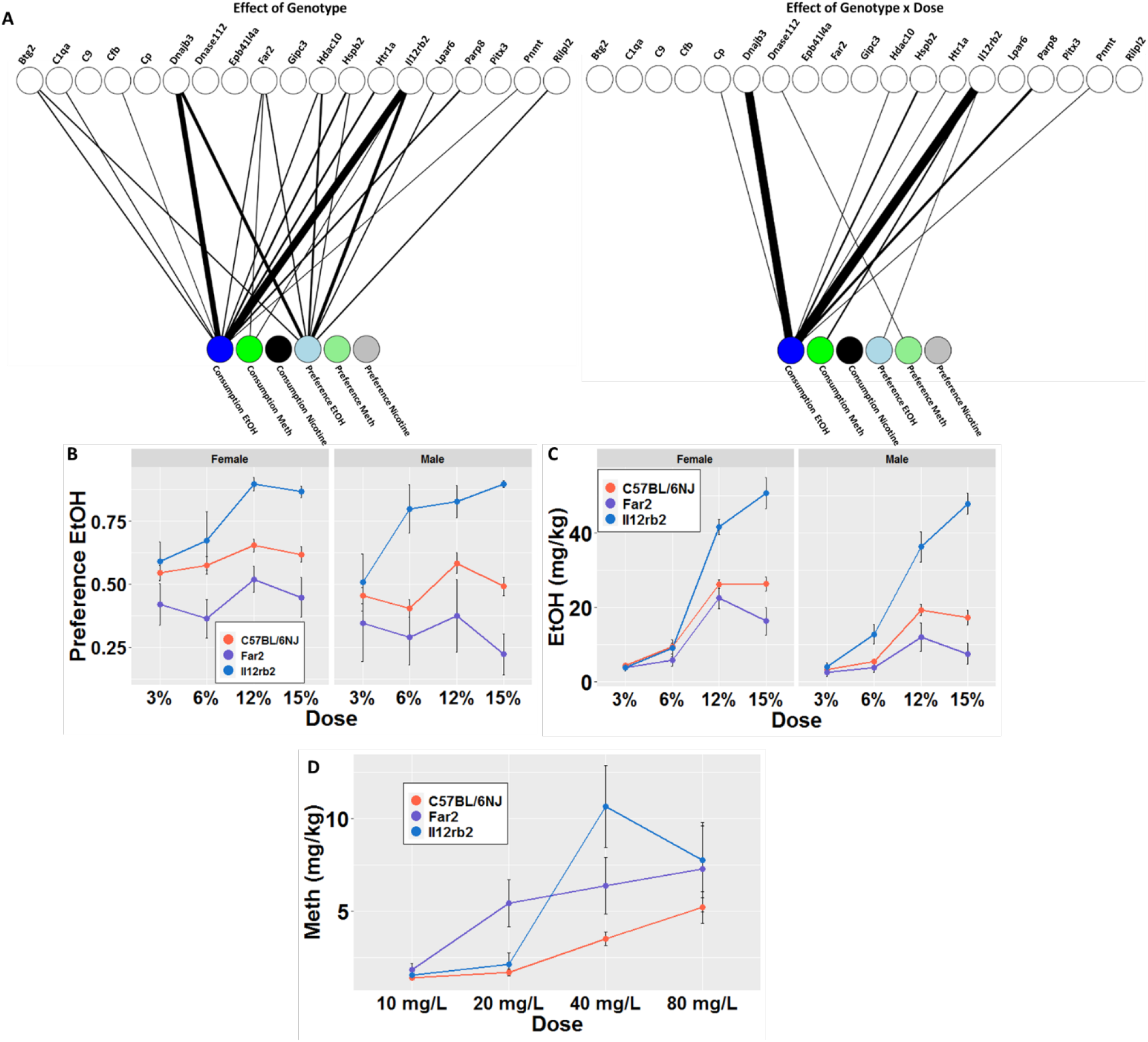
Shared associations of gene deletion mutations with alcohol, methamphetamine, and nicotine use phenotypes. A) Bipartite graphs depicting significant effects of strain and strain × dose for consumption and preference of alcohol (EtOH), nicotine, and methamphetamine (MA). Significant associations are represented by the thickness of the edge connecting the two nodes. Edge weights are inversely proportional to the –log10 p-value of the association. B) Dose-response curve depicting the effect of *Il12rb* and *Far2* deletions on EtOH preference. C) Dose-response curve depicting the effect of *Il12rb* and *Far2* deletions on EtOH consumption. D) Dose-response curves depicting the effect of *Il12rb* and *Far2* deletion on MA consumption. Data are shown as mean ± SD (FDR < 0.05).

Oral EtOH-use phenotypes (consumption and preference) showed the highest numbers of significant associations; five strains showed overall strain effects, and eight showed strain × dose effects for at least one of the two EtOH phenotypes (Fig. 2A). We further found a wide range of outcomes exemplified by *Il12rb2* and *Far2*. Deletion of *Il12rb2* resulted in an increase in both EtOH preference (F_strain (1, 90)_ = 35.7, FDR= 8.63E-07) and consumption (F_strain (1, 90)_ = 88.48, FDR=9.28E-14) compared to C57BL/6NJ controls (Fig. 2B). In contrast, deletion of *Far2* resulted in a decrease in both EtOH preference (F_strain (1,87)_ = 12.31, FDR=3.40E-3) and consumption (F_strain(1,87)_ = 9.3, FDR=6.40E-3) as compared to controls (Fig. 2C). Across the strains, females (F_sex (1, 318)_ = 18.16, P= 2.68E-05) exhibited higher levels of preference and consumption for EtOH than males (F_sex (1, 318)_ = 34.03, P= 1.34E-08).

For MA, in which we tested 17 of the 19 single-gene KO strains, we found that three strains (*Il12rb2*, *Far2,* and *Dnase1l2*) showed significantly altered preference or consumption of MA (Fig. 2A). Unlike for EtOH outcomes, we did not detect sex effects for MA phenotypes. Interestingly, while *Il12rb2* and *Far2* deletions showed increased and decreased responses, respectively, for oral EtOH self-administration phenotypes compared to controls (Fig. 2D), *Il12rb2* and *Far2* deletions both led to significantly increased consumption of MA (*Il12rb2*, F_strain (1, 71)_ = 9.88, FDR=2.19E-02; Far2, F_strain (1, 69)_= 10.59, FDR=2.19E-02) (Fig. 2C).

Using the 2BC screening, we could not detect any KO strains with significantly altered oral nicotine self-administration phenotypes (Fig. 2A). These findings could be due to the aversive taste of oral nicotine, a confounding effect of saccharine with the nicotine, or a reflection of the small genetic effect size for the measured phenotypes most likely due to the complexity of the nicotine’s pharmacology both in terms of dose-response but also temporal patterns^3,54,55^.

### 3.3 Principal Component Analysis to Define Relationships Among Phenotypes

PCA revealed relationships within and between the ten-predisposing drug-naïve phenotypes, six drug self-administration traits (consumption and preference × three drugs), and six liquid intake traits (water-drinking and total fluid intake × three drugs), a total of 22 measured traits (Table S5). We included liquid intake traits in this analysis to account for variation in total fluid consumption unrelated to the drug. We then performed bi-plot clustering using the effect sizes across all 22 traits for the 16 KO strains tested for all three drugs (Figure 3). ^56^. The PCA reveals that principal components one and two account for 21.1% and 17.8% of the variance respectively, together accounting for ∼39% of the variation observed in our 16 strains across the 22 measures. PC1 differentiates KO strains with ethanol drinking from those that display non-ethanol drinking phenotypes. PC2 relates ethanol-drinking KO strains to drug-naïve behavioral profiles of high or low exploration.

**Figure 3.**
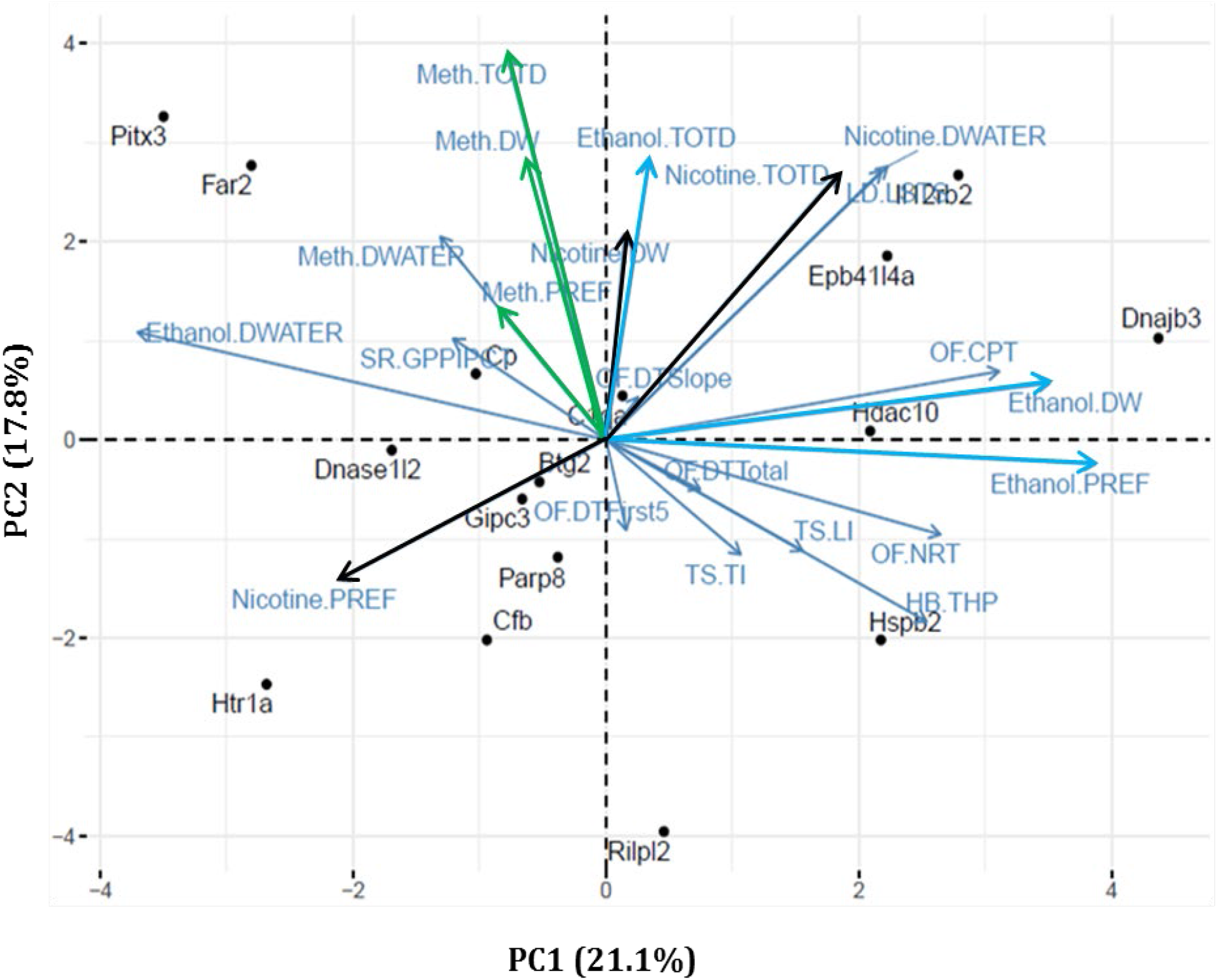
Shared relationships among predisposing drug-naïve behaviors with drug-use and liquid intake phenotypes. Principal components analysis was used to assess shared variance among predisposing drug-naive behavioral traits and drug-use and liquid intake phenotypes. Each point represents a KO strain, while arrows represent each of the analyzed traits. Analysis was conducted on all 16 strains tested on all behavioral phenotyping measures. (Colors are added to drug-use phenotypes to match color scheme from Figure 2, Nicotine is black, dark blue is EtOH and green is MA. Light blue refers to water consumption or base line behaviors) HB: Hole board, THP: Total hole pokes, LD-LSTS: light/dark time spent in light, OF-CPT: Center Permanence Time, OF-DTFirst5: Distance Traveled First Five Minutes, OF-DTSlope: Distance Traveled Slope, OF-DTTotal: Distance Traveled Total, OF-NRT: Number of Rears Total, SR-GPPIPCT: Amplitude Percent PPI Global, TS-LI: Latency to immobility, TS-TI: Time immobile, Ethanol-DW: Ethanol Consumed, Meth-DW: Meth Consumed, Nicotine-DW: Nicotine Consumed, Ethanol-PREF: Preference Ethanol, Meth-PREF: Preference Meth, Nicotine-PREF: Preference Nicotine, Ethanol-TOTD: Total Drinking Ethanol, Meth-TOTD: Total Drinking Meth, Nicotine-TOTD: Total Drinking Nicotine, Ethanol-DWATER: Water Drinking Ethanol, Meth-DWATER: Water Drinking Meth, Nicotine-DWATER: Water Drinking Nicotine

The correlations within and between any of the predisposing drug-naïve phenotypes and drug-use phenotypes can be assessed using the angle which separates any two vectors (Fig. 3). Vectors that fall close to one another (where the angle approaches 0°) are strongly positively correlated; vectors which fall 180° apart are strongly negatively correlated; and vectors which fall 90° apart are independent of one another^57^. Using these relationships, we can use the PC1 axis to classify our tested strains. For example, we can separate which strains had increased EtOH consuming/preferring phenotypes from those with decreased EtOH consuming/preferring phenotypes. Along the PC2 axis, we found clustering of predisposing drug-naïve phenotypes, which can be used to divide our strains into different baseline behavioral profiles, i.e., “low anxiety” or “exploratory” profiles. Also, along PC2, we see a close relationship that separates our MA-consuming strains from our non-consuming strains. Scores for each strain are obtained by multiplying the PC loadings by the strain means, allowing strains to be plotted in the two-dimensional space. Strains with high absolute scores on both ends of PC1, such as *Il12rb2* and *Far2,* have strong opposing increased and decreased EtOH preferring/consumption phenotypes, respectively. Additionally, *Il12rb2* and *Far2,* whose variation is similarly explained by PC2, show that while these two strains manifest opposing EtOH phenotypes, both manifest strongly increased MA consumption phenotypes.

### 3.4 Phenotypic Deviance Based on Mahalanobis Distance for Predisposing Drug-Naïve Phenotypes in Cohort #2

We calculated overall phenodeviance using Mahalanobis distances^33^ of effect sizes relative to C57BL/6NJ controls. Using this calculation, we represented the phenodeviance from the C57BL/6NJ controls across all eight measured predisposing drug-naïve phenotypes as a single score (Fig. 4). Higher scores represent greater overall phenodeviance from controls across all predisposing drug-naïve phenotypes. In 2017, 401 strains had completed the phenotyping pipeline and were included in the analysis. Results from this analysis indicate that of the 401 single-gene KO strains tested, 123 strains were significantly phenodeviant, with scores ranging from 24.1-2038.9. This range suggests that even within the significantly phenodeviant strains, strains with exponentially greater predisposing drug-naïve phenotypes and risk factors existed, making them the most likely to manifest drug-use phenotypes and potentially have multi-drug effects. Therefore, we prioritized the top 25 strains with high Mahalanobis scores (≥517.3) that were also available for rederivation or could be directly obtained to screen for drug-use phenotypes. Of these, fifteen significantly phenodeviant KO strains were successfully rederived or obtained and bred for testing, thirteen of which scored in the top quartile of Mahalanobis scores (Fig. 4). We also included the *Tmod2* and *Rap2b* KO strains, which were phenodeviant as determined by the Mahalanobis distance calculations but not in the top 25 available strains that were most phenodeviant.

**Figure 4:**
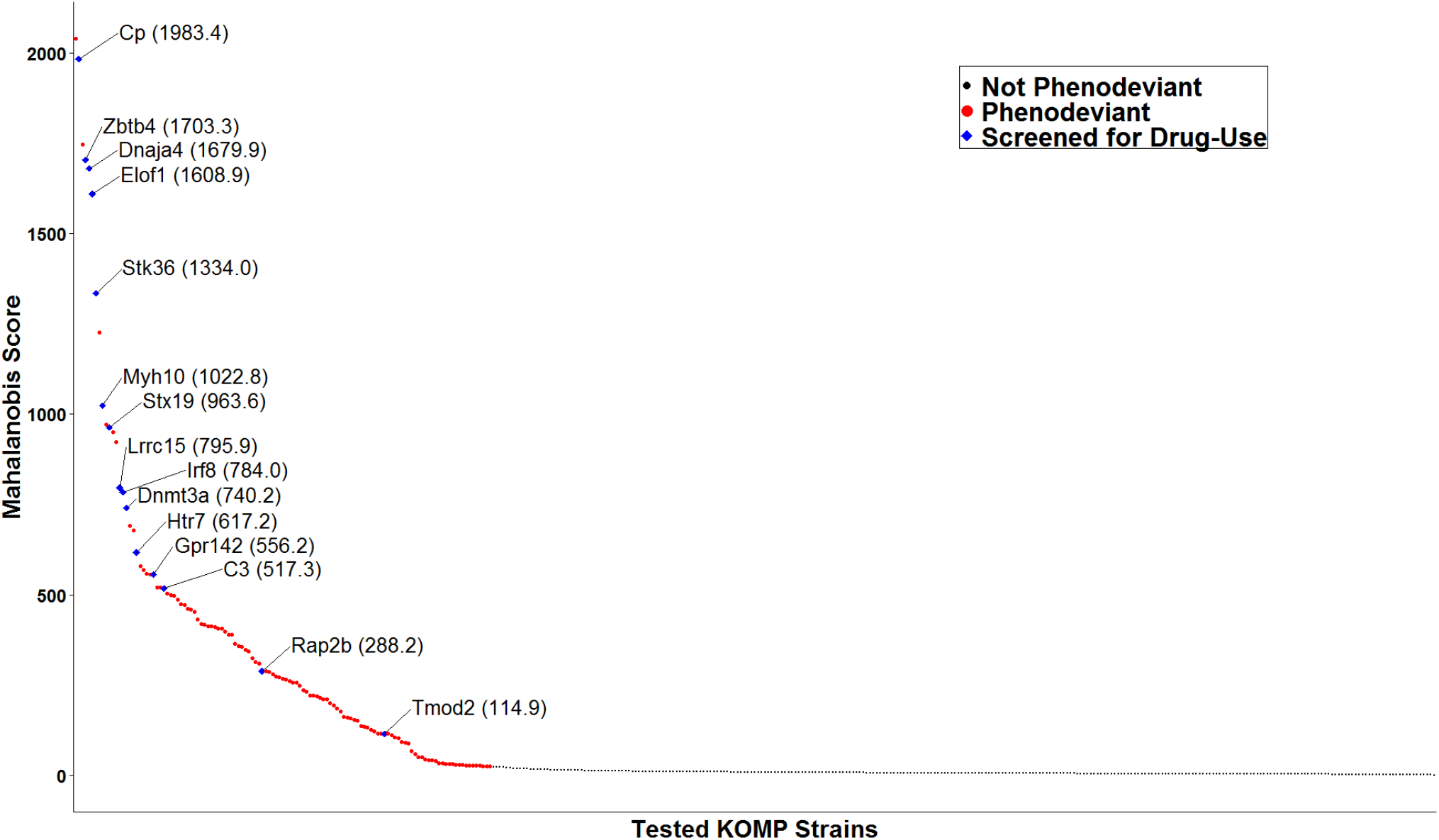
Multidimensional assessment of phenodeviance in drug-naïve behaviors. Of the 401 strains tested, the 123 KO strains, each indicated as a colored circle in the plot, showed a statistically significant difference from matched C57BL6/NJ controls using Mahalanobis score (FDR < 0.05). Red circles represent single gene KO strains that were not rederived; blue points represent strains that were rederived. All blue points are identified by their gene abbreviations and Mahalanobis scores.

#### 3.4.1 Two-Bottle Choice Assay to Determine Methamphetamine-Use Phenotypes

In our initial screening of the KO strains tested by the MA 2BC assay in Cohort #1, three of the 19 strains (*Il12rb2*, *Far2,* and *Dnase1l2*) had altered consumption or preference attributed to the main effects of strain or strain × dose (FDR<0.05). Sex had no significant effect on either MA consumption or preference phenotypes. Of the 15 strains identified as phenodeviant by Malanhobis testing in Cohort #2, eight strains had significant MA preference phenotypes revealed by bipartite analysis (Fig. 5A), either manifested through a main effect of strain or strain × dose (FDR < 0.05). Of these eight strains with MA preference phenotypes, only one (*Cp*) was also present in the 19 phenodeviant strains identified by PCA in Cohort #1. Focusing on strains that showed phenotypes in multiple drugs, strains *Irf8, Tmod2,* and *Rap2b* all exhibited significantly increased preference for MA (F_strain (1, 51)_ = 12.34, FDR= 4.70E-03), (F_strain (1, 51)_ = 15.23, FDR= 2.10E-03), and (F_strain (1, 51)_ = 15.14, FDR= 4.37E-03), respectively, compared to the control strain (Fig 5B). The *Irf8* strain showed a significant interaction of strain × dose and exhibited significantly increased preference to initial lower doses of 10 mg/L and 20 mg/L increasing to 55.1 ±6.5% and 40.0 ±6.4%, from 31.51 ±3.3% and 26.64% ±3.0% respectively and did not display differences from control strains at higher doses (F_strain × dose (3, 140)_ = 3.69, FDR= 4.05E-02). Additionally, the *Irf8, Tmod2,* and*Rap2b* (not depicted) strains consumed more MA over the 2BC protocol than control strains doses (F_strain (1, 51)_ = 7.42, FDR= 4.40E-02 and F_strain (1, 51)_ = 8.92, FDR= 3.24E-02, respectively) (Fig. 5C).

**Figure 5.**
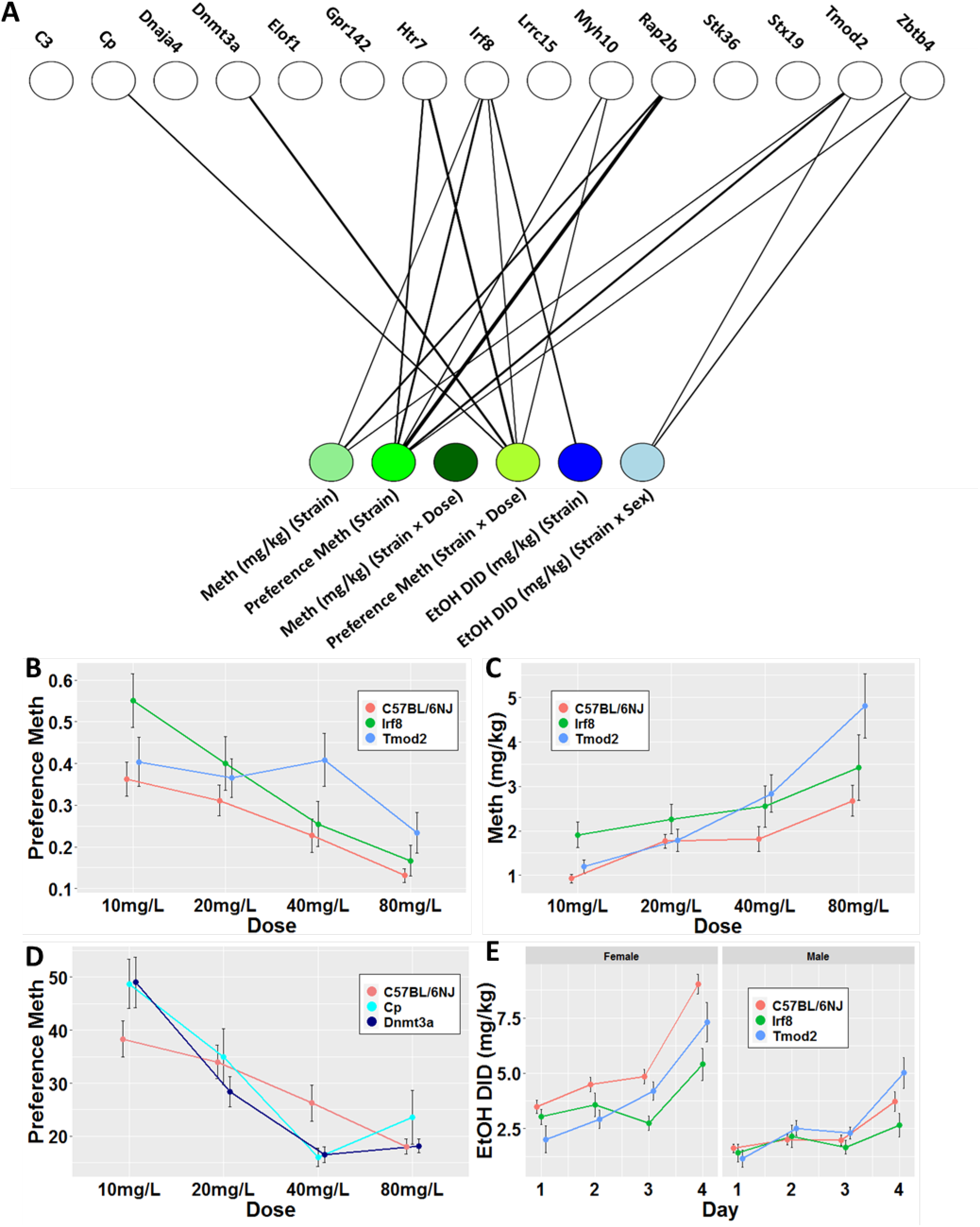
Single-gene KOs resulting in significant drug-use effects cohort 2. A) Bipartite graph displays significant hits across the three measured phenotypes. Graph depicts significant effect of strain with the three green nodes and effects of strain by dose with the three blue nodes. Significant associations are represented by the thickness of the edge connecting the two nodes. Edge weights are inversely proportional to the –log10 p-value of the association. B) Dose-response curve depicting the effect of *Irf8* and *Tmod2* deletions on MA preference. C) Dose-response curve depicting the effect of *Irf8* and *Tmod2* deletions on MA consumption. Data are shown as mean ± SD for n = 16 in each KO group. D) Dose-response curve depicting the effect of *Cp* and *Dnmt3a* deletions on MA preference. E) Dose-response curves depicting the effect of *Irf8* and *Tmod2* deletion on EtOH DID consumption. Data are shown as mean ± SD for n = 16 in each KO group.

For the control and most KO strains evaluated, preference for MA trended down as doses increased, with 10 mg/L being the dose with the highest preference compared to water. The control strain had its highest average MA preference, 36.25% ± 4.0%, for the initial 10 mg/L dose (Fig. 5B) and consumed higher percentages of water at all doses. At the initial 10 mg/L dose, *Irf8* consumed a higher percentage of MA, 55.1±6.5%, than water. In general, *Irf8* had a higher preference for MA than controls, but differences were largely exhibited in the initial two doses (Fig. 5B). While most strains had their greatest preference for MA at the lowest dose of 10 mg/L similar to controls, we observed a shift in the dose-response curve for *Tmod2.* The *Tmod2* strain had similar preference levels to the initial dose as controls but had a peak for MA preference at the 40 mg/L dose (40.9±6.4%) (Fig. 5B). Further studies will be needed to determine to what drives the increased preference for sensitivity to methamphetamine^58^.

In the second screening, two strains with deletion of genes with distinct biological functions (*Dnmt3a* and *Cp*) resulted in very similar alterations to MA preference in a dose-dependent manner. Initially, both *Dnmt3a* and *Cp* exhibited increased preference for MA at the initial dose compared to controls (49.1±5.6% and 49.0±5.4%, respectively). We did not observe any differences from controls at 20mg/L dose. However, at the higher dose of 40 mg/L, both strains showed a decreased preference for MA compared to controls (9.7±1.3% for *Dnmt3a* and 10.2±2.0% for *Cp* vs.17.1±2.7% for controls). These results suggest that the deletion of these two genes alters sensitivity to MA. While the initial preference was non-aversive and equal to the percentage of water consumed, it became more rapidly aversive as doses increased compared to controls (Fig. 5D).

#### 3.4.2 Drinking in the Dark to Determine Ethanol-Use Phenotypes

Analysis of DID results indicate that three of the 15 phenodeviant strains in Cohort #2 displayed significantly altered EtOH consumption (*Irf8, Tmod2,* and *Zbt64*). EtOH DID consumption was strongly influenced by sex in all strains (F_sex (1, 297)_ = 271.3, p < 2.22E-16). *Irf8* (Fig 5E) had significantly altered EtOH DID consumption across the four days of access (F_strain (1, 48)_ = 13.38, FDR= 9.48E-03). Additionally, for two strains, *Tmod2* (Fig 5E) and *Zbtb4* (not depicted), EtOH DID consumption was influenced by strain and sex (F _strain × sex (1, 47)_ = 8.39, FDR= 4.27E-02) and (F_strain × sex (1, 48)_ = 11.72, FDR= 1.91E-02), respectively. Similar to what was observed in the EtOH DID consumption phenotype, BEC was also influenced by sex in all strains (F_sex (1, 264)_ = 33.01, P= 2.52E-08). Of the 15 phenodeviant strains chosen for testing in the DID paradigm, no strain resulted in significantly different BEC from the control strain.

#### 3.4.3 Genes with Drug-Specific or Multi-Drug Effects

Using a multidimensional assessment of phenodeviance across predisposing drug-naïve phenotypes in Cohort #2, we identified 15 single-gene KO strains that we tested in a second screen for altered patterns of drug-use phenotypes. Results from MA 2BC from the second cohort in combinations with EtOH DID revealed that eight out of 15 (53.3%) of our identified single-gene deletions resulted in an altered drug self-administration phenotype. Three (38%) of these strains had multi-drug effects across both drugs (Fig 5A). Importantly, the second screening process and ability to rederive more extreme phenodeviant strains increased our ability to identify genes with multi-drug effects from 12.5% (2/16) to 20% (3/15). However, our overall hit rate was higher for individual drugs in the original cohort tested for which 15 gene deletions that showed both predisposing drug-naïve phenotypes and drug self-administration phenotypes in Cohort #1 (79%, 15/19).

#### 3.4.4 Functional Analysis of Candidate Genes Reveals Diverse Mechanisms of Involvement in Addiction Related Phenotypes

Although few if any of the genes we evaluated were recognized as addiction related genes in the literature at the time of testing, 22 genes we identified with both predisposing drug-naïve phenotypes and drug-use phenotypes were supported by additional evidence from at least one of the searched databases establishing a prior connection to drug-related studies, either through expression data, QTL mapping, or connections to drug-related biological mechanisms (Table S6). In addition to the functional analysis of drug-related gene sets in GeneWeaver, we assessed the genes with significant effects for overlapping representation in biological pathways. Using GSEA, a systematic search of Canonical, KEGG, and GO biological or cellular pathways revealed that none of our genes were annotated to the same pathways. (Accession date 02/04/2020 & 05/08/20). These results suggest that each of the single-gene KOs may all alter drug-use phenotypes through multiple independent biological pathways and mechanisms.

## DISCUSSION

Overall, our data indicate the utility of leveraging the known complex relationships among predisposing drug-naïve phenotypes and their drug-related addiction risk phenotypes. In this project, we used and refined our understanding of these relationships in combination with the high-throughput JAX-KOMP2 program to identify thirty-three plausible single-gene KO strains predictive of drug-use phenotypes. Of those thirty-three plausible candidates, 22 (67%) of the single gene KOs significantly altered at least one drug-use phenotype. Following screening for drug-use phenotypes, we validated all significant genes through functional analysis for plausible connections and/or mechanisms through which they potentially could have altered drug-use phenotypes. Further analysis through GSEA indicated no overlapping pathways among our candidate genes that could have possibly affected drug-use phenotypes, suggesting that these novel candidate genes could represent multiple diverse pathways for roles in drug use.

The strategy we used to identify drug-use candidate genes using predisposing drug-naïve phenotypes was successful and circumvented the effects of drug exposure on subsequent physiological testing in the screening program, allowing us to discriminate risk from consequences of drug exposure. An approach that uses a drug-naïve screen is efficient, but it will necessarily miss those genes with drug-use effects that are not manifested in predisposing drug-naïve phenotypes. Nevertheless, through this study, combined with publicly available data, multiple novel candidate genes, high-throughput testing using multiple drugs, and functional analysis of multiple genomic databases, we have identified 22 new drug-use genes amenable for detailed characterization in viable mutant mice (Table S6).

The initial screening identified 15 novel drug-use gene candidates leveraging data from predisposing drug-naïve phenotypes, corroborating previous studies that found shared genetic components underlying predisposing drug-naïve phenotypes and subsequently drug-use phenotypes^20,21^. Interestingly, in contrast to findings in the literature^8,59-61^, the relationships we found were not uniform connections between drug-use phenotypes and their predisposing drug-naïve phenotypes. Our results indicated more complex and multidimensional relationships that we analyzed further using PCA. In this analysis, strains with significant drug-use phenotypes were found in all four quadrants of the graph (Fig 3), each representing a different baseline behavioral profile predictive of different drug phenotypes. Two ethanol-preferring strains that exemplify different baseline behavioral profiles were *Il12rb2* (found among strains with risk-taking/low-avoidance behaviors) and *Hspb2* (found among strains with high exploratory/ high activity behaviors). Although both KO strains showed an ethanol-preferring phenotype, the different behavioral profiles segregated along PC2, also correlating with an MA consumption phenotype. The 2BC choice data reveals that *Il12rb2* KO mice have a significant MA consumption phenotype, whereas *Hspb2* KO mice do not. Thus, results from the PCA revealed the diverse multidimensional nature of the relations underlying the many predisposing behaviors and their predicted drug-use phenotypes. Rather than reflecting a uniform predictive relationship between each behavioral phenotype and its predisposing effect on drug intake^8,59-61^, these findings indicate a complex interaction of all the predisposing behaviors and their effects on drug-use phenotypes across different drugs, and that many biological mechanisms support the distinct relations among baseline behaviors and drug-use phenotypes. They corroborate and extend to psychostimulants, the previous work of Blednov and colleagues which indicated that distinct mutations, albeit on heterogeneous backgrounds, disrupt multiple physiological systems associated with ethanol consumption.^62^ The lack of overlapping pathway membership observed for the detected genes further reveals the tremendous breadth of variation that can result in addiction-related phenotypes and the potential for sizeable individual variation in mechanisms of addiction vulnerability among those with SUD. Through deeper exploration of these relationships, we can better understand the specific relationships among biological pathways and behavioral processes that lead to heterogeneous behavioral and genetic mechanisms of addiction and substance use.

Much of the historical focus in addiction research has been on studying genetic components underlying drug-specific effects through alteration of drug-specific metabolism or drug receptors in the reward pathway^20,63-67^. These genetic components can play crucial roles in the development of treatments for drug-specific SUDs. Interestingly, a functional analysis using GeneWeaver and GSEA revealed that the thirteen genes have diverse functions and expression patterns with no annotated pathway overlap or any enrichment for similar GO terminology. These results suggest that these genes may each represent independent biological pathways and mechanisms involved in vulnerability to EtOH use and warrant further characterization. For example, *Il12rb2* and *Far2*, the two genes that showed multi-drug effects (i.e., significant alteration to both EtOH and MA), have diverse biological functions and expression patterns and no enrichment for similar GO terms. *Il12rb2* (interleukin 12 receptor subunit beta 2) is a subunit of the interleukin 12 receptor complex involved in IL12-dependent signaling and functions in Th1 cell differentiation. It is highly expressed in the pancreas, placenta, skeletal muscle, NK cells, and multiple brain regions. In contrast, *Far2* (fatty acyl-CoA reductase 2) is a member of the short-chain dehydrogenase/reductase superfamily that functions in fatty acid metabolism. It is highly expressed in intestinal tissue, white blood cells, epididymis, and multiple brain regions.

We identified 22 genes not previously connected to drug use, which significantly affected both predisposing drug-naïve and drug-self-administration phenotypes when knocked out. Functional analysis of these genes revealed that the only significant overlap was between *Htr1a* and *Htr7*, which are both part of the canonical pathway for REACTOME_SEROTONIN_RECEPTORS (M6034). Additionally, an extensive literature search revealed direct connections between *Il12rb2* and *Irf8* as part of the cytokine-mediated pro-inflammatory immune response of the central nervous system. The few numbers of connections observed between identified genes suggest that, for the most part, all these novel gene candidates potentially represent distinct mechanisms for drug-use vulnerability.

Although our primary goal was to elucidate gene-specific effects on predisposing addiction behaviors, we were also interested in the interaction of genotype and sex. Our results corroborate findings from previous studies, which found that sex differences did not have significant effects on MA use^65,67^ but did significantly influence EtOH-use^68-70^, with female mice consuming higher levels of EtOH than male mice. The only significant strain × sex interaction we observed was in our EtOH DID protocol, where *Tmod2* and *Zbtb4* had significant strain × sex interactions as indicated by decreased consumption for the female strains but no difference in male consumption compared to controls. These results suggest that these genes could be differentially regulated in each sex and their deletion results in more similar drug-use phenotypes between the sexes. The findings of strain × sex differences in responses of *Tmod2* and *Zbtb4* to EtOH are particularly interesting because these two strains also showed significantly altered MA intake but no effect of sex or strain × sex. Additionally for *Tmod2*, previous studies conducted using strains from the BXD recombinant inbred mice strains found sex differences in gene expression in various locations throughout the reward pathway following drug exposure^19^. Together, these findings suggest that these genes could potentially be regulated in a strain × sex × drug manner.

Addiction is a multi-phased process, and the genetic mechanisms associated with sustained drug-use may be independent from that of the transition from initial use to addiction. Further characterization of genes involved in addiction-related behavior and associated pathways could elucidate their distinct roles in the process of transition addiction. Our findings suggest that evaluating single-gene KO mice using a broad neurobehavioral screen allows the continued identification of novel addiction risk genes. In this project, we detected multiple genes affecting drug-use phenotypes through diverse biological pathways. Of the many diverse pathways represented by our identified drug-use genes, we highlighted the potential role of the neuroimmune and cytokine responses in altering drug use which connected three of our novel drug-use genes with the strongest effects across drugs. Each of these genes would only account for small proportions of the genetic variation and would often be missed using GWAS. The continual screening of KO mice for predisposing drug-naïve phenotypes can lead to the discovery of previously undetected addiction risk genes across the breadth of pathways involved in these devastating conditions.

## Acknowledgements

This work was supported by UM OD 023222, and a supplement funded by NIH CRAN (National Institute of Drug Abuse and National Institute of Alcohol Abuse and Alcoholism, National Cancer Institute), and DA039841 Center for Systems Neurogenetics of Addiction. PED was funded by NIDA K99 DA043573 and JAB by NIDA U01 DA043809 during preparation of this manuscript. Stephen Krasinski provided substantial editing assistance during preparation of the manuscript. We are extremely grateful to the technical work performed by the members of the JAX KOMP mouse production and mouse phenotyping teams without whom this work would not have been possible, Scientific services and CCSG P30 CA034196 for support of V.M.P and others in the Computational Sciences Service, the Center for Biometric Analysis and Clinical Chemistry.

