## Supplemental Materials for "DISCOVERY AND VALIDATION OF GENES DRIVING DRUG-INTAKE AND RELATED BEHAVIORAL TRAITS IN MICE"

Supplementary Tables

Supplementary Figure 1: Behavioral tests used to select KOs with potential drug-use phenotypes from JAX KOMP2 Pipeline.

Supplementary Table 1: KOMP2 Pipeline

Supplementary Table 2: Strain sample sizes by drug for first round of two bottle choice screening.

Supplementary Table 3: Strain sample sizes for second round of methamphetamine two bottle choice screening.

Supplementary Table 4: Strain sample sizes for ethanol drinking in the dark from second screening

Supplementary Table 5: Factor Score Loadings to Principal Components

Supplementary Table 6: Summary of results from bioinformatics search connecting significant knockout genes to other drug/ addiction related studies.

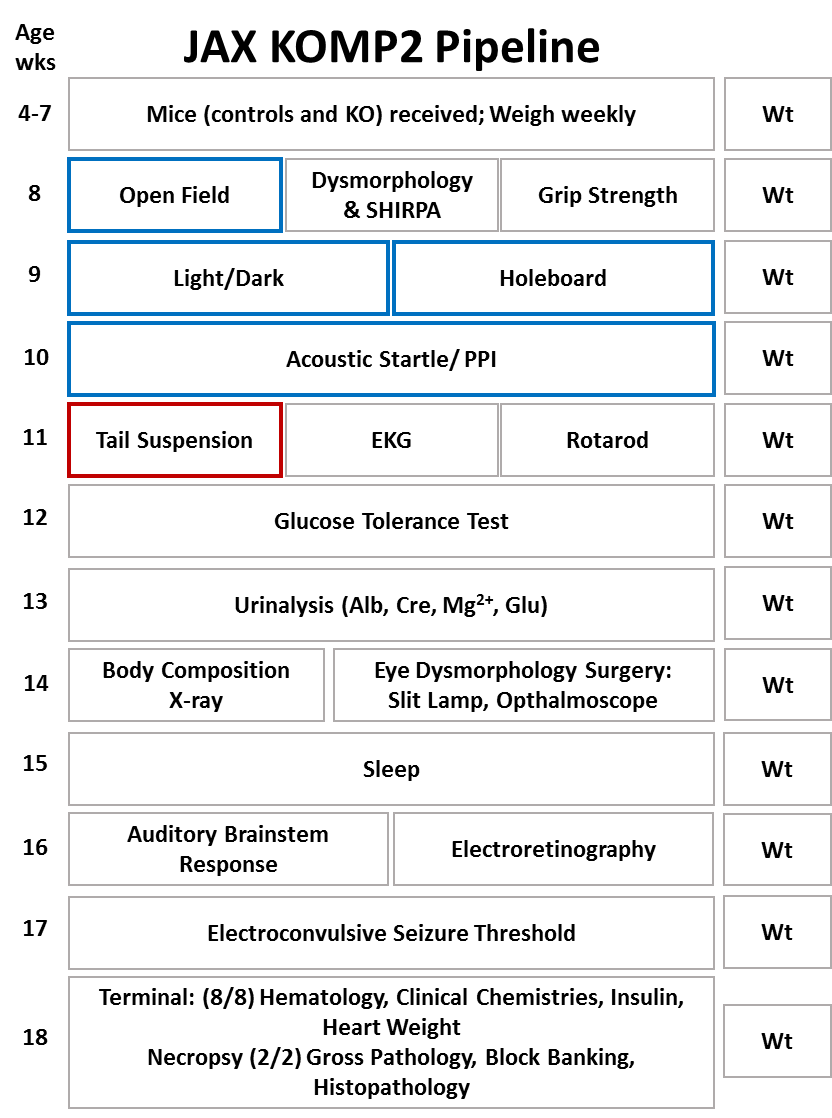

**Supplemental Figure 1: Phenotyping pipeline of JAX KOMP2 Project.** All specific phenotyping analysis by age is shown for the JAX KOMP2 Project. Data from the behavioral tests outlined in blue and red were used to identify KO strains with predisposing phenotypes in the first cohort (2014). Data from the behavioral test outlined in blue was used to identify KO strains with predisposing phenotypes in the second cohort (2017) because tail suspension had been discontinued.

**Supplementary Table 1: Rationale for predisposing drug-naïve phenotypes***

| **Test** | **Phenotype** | **Rationale** |
| --- | --- | --- |
| **Open Field** | **Total Rearing** | The total amount of times a mouse rears or jumps during the twenty-minute testing session in the open field arena. This endpoint serves as a measure for anxiety^1,2^ which is positively genetically correlated with substance use in mice.^3^ Decreased total rearing is an index for increased anxiety. |
|  | **Total Distance Travelled** | Total distance traveled during the twenty-minute testing session in the open field arena. Distance traveled serves as a measure of anxiety ^1,2^ which is positively genetically correlated with substance use in mice.^3^ Decreased total distance travelled is an index of increased anxiety. |
|  | **Slope of Distance Travelled** | Slope of the best fit line measuring the change in distance traveled over the twenty-minute test which is broken into four five-minute time bins. This measure is calculated to show habituation to the novelty which can be used to measure anxiety or exploratory/risk-taking phenotypes.^4^ These behaviors are genetically correlated with substance use in mice. ^3^ Increased slope (increased habituation) is an index for decreased anxiety. |
|  | **Distance First Five Minutes** | Total distance traveled during the first five minutes of the 20-minute test in the open arena. Distance traveled in the first five minutes of open field serves as a measure of novelty reactivity and anxiety^5^ which has been shown to be predictive of initiation of drug use and progression to compulsive drug use.^6^ High distance travelled reflects high novelty reactivity and low anxiety. |
|  | **Center Time** | The total amount of time spent in the center 40% of total surface area in the open arena during 20-minute testing. Time spent in the center servers as a measure for anxiety^1,6^ and is positively genetically correlated with substance use in mice.^3^ High time in the center reflects low anxiety. |
| **Light/Dark** | **Time in Light** | The total amount of time, represented as a percentage of total testing time, during which the mouse spent on the light side of the two-chambered light dark apparatus during the single 20-minute testing session. Time in light is an index of anxiety^7^ and is positively genetically correlated with substance use in mice.^3^ Less time in light relative to controlsindicates high anxiety. More time in light than controls reflects excessive risk-taking, a form of impulsivity. |
| **Holeboard** | **Holepokes** | The total number of nose-pokes into the 16 holes in the hole board testing arena during the single 20-minute testing session. Nose-pokes in a hole board is one of several genetically distinct indexes of novelty seeking^8^ which are positive predictors of substance use in mice.^9^ High nose-pokes reflect high novelty seeking. |
| **Acoustic Startle** | **Percent Prepulse Inhibition (%PPI)** | Percentage of baseline startle response when lower-intensity ‘prepulse’ sounds precede a louder ‘pulse’ sound. Reduced PPI is a Research Diagnostic Criteria (RDOC) and endophenotype for multiple neuropsychiatric disorders^10^ including panic disorder (anxiety) which is positively genetically correlated with substance use in mice ^3^. Reduced %PPI is an index for increased anxiety. |
| **Tail Suspension** | **Time Immobile** | The total amount of time a mouse is immobile while suspended by its tail during the five-minute testing session. Time immobile is recognized as an animal model for efficacy of antidepressants^11^ which has been shown to increase chances of drug acquisition and maintenance.^12^ While there may be controversy over the validity of this model as a depression-like behavior, many researchers still use time immobile due to its predictive validity.^13,14^ A greater time immobile is considered a measure of higher levels of depression-like behavior. |
|  | **Latency to Immobility** | The total amount of time a mouse is actively moving while suspended by its tail before it becomes immobile during the five-minute testing session. Latency to immobility is recognized as an animal model for the study of depression^11^ which has been shown to predict increased drug acquisition and maintenance.^12^ Shorter latency to immobility reflects greater depression-like behavior. |

*****The table depicts the 10 behavioral traits chosen from the five behavioral tests within the KOMP2 pipeline. Additionally, the table explains what each of the ten chosen behavioral phenotypes measure and why they are viewed as predisposing drug-naïve phenotypes.

**Table S2: Strain sample sizes by drug for first round of two bottle choice screening.**

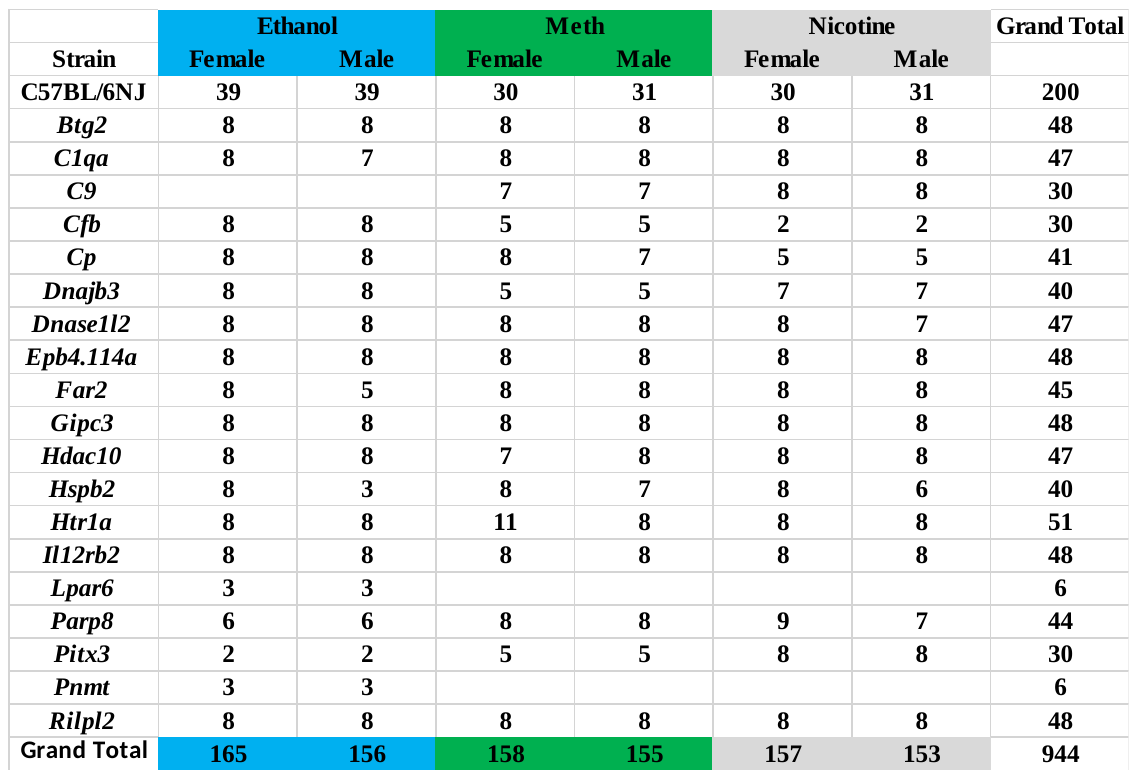

**Table S3: Strain sample sizes for second round of methamphetamine two bottle choice screening.**

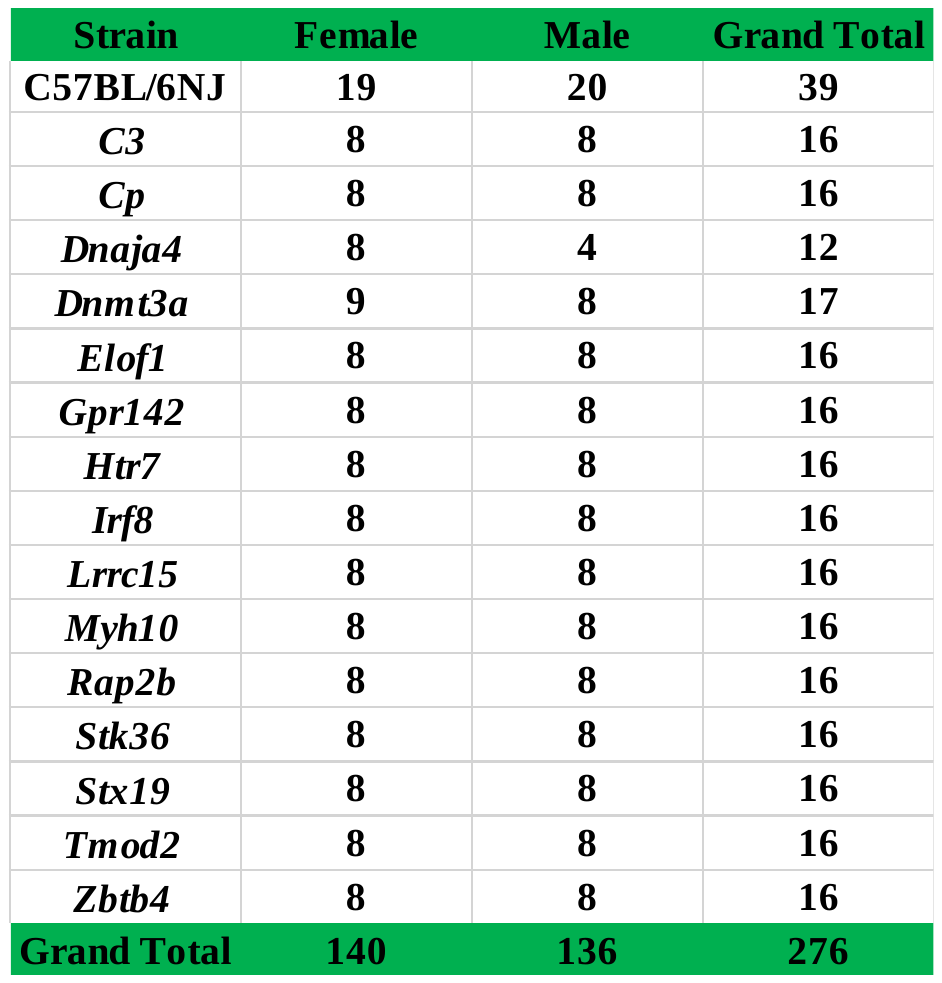

**Table S4: Strain sample sizes for ethanol drinking in the dark from second screening**

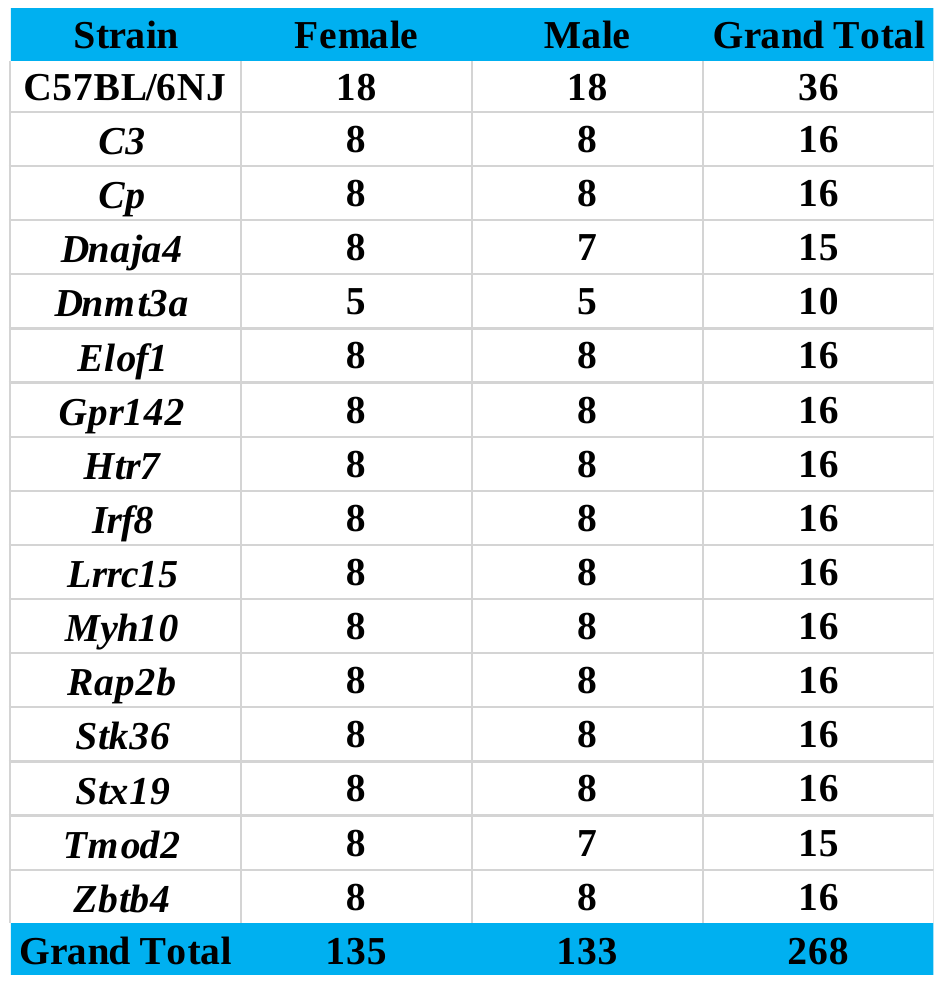

**Table S5: Factor score loadings to principal components**

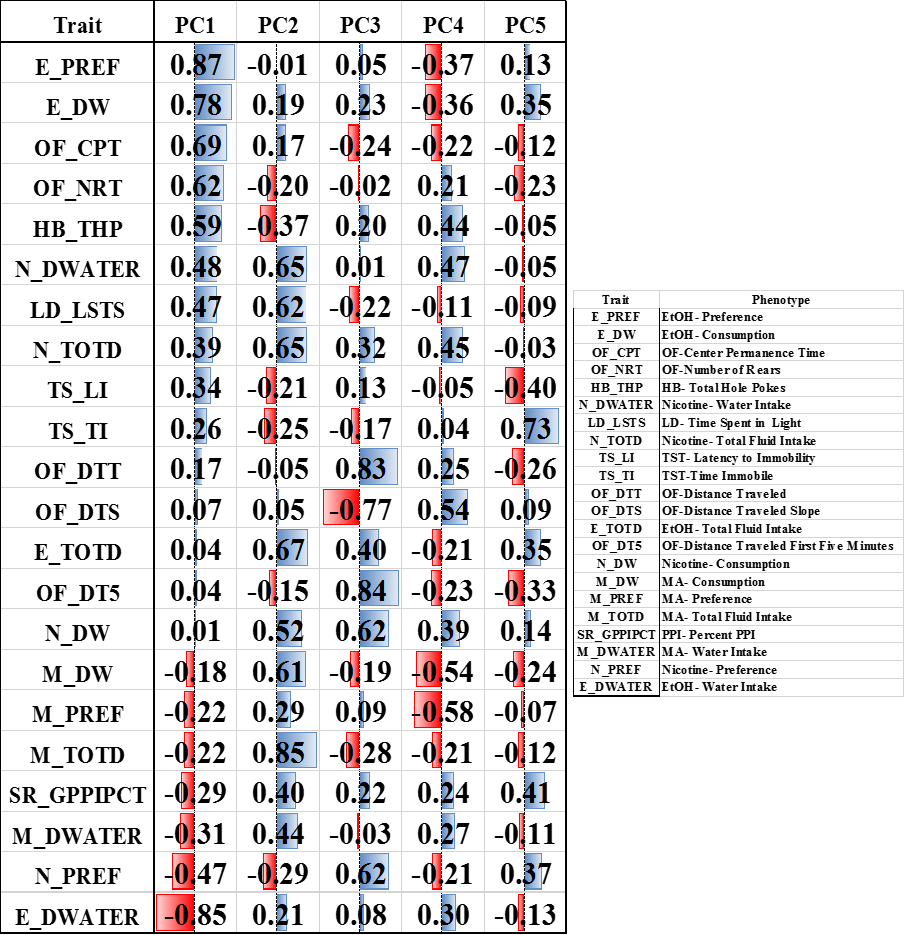

**Table S6: Summary of results from bioinformatics search connecting significant knockout genes to other drug/ addiction related studies**

| **Gene** | **Geneset ID** | **Database** | **Description of Geneset** |
| --- | --- | --- | --- |
| Hdac10 | GS233344 | KEGG Geneset | "Alcoholism" pathway genes, |
|  | GS233499 | KEGG Geneset | "Alcoholism" pathway genes |
|  | GS233931 | KEGG Geneset | "Alcoholism" pathway genes |
|  | GS86789 | [DRG] | Table S1: Cocaine Regulation of H3 Acetylation. (provisional) |
| Lpar6 | GS86977 | [DRG] | Table S1: All transcripts significantly different in abundance between the majority of heroin subjects and their matched controls (provisional) |
|  | GS84277 | (Published QTL ) | METH responses for home cage activity |
|  | GS84278 | (Published QTL ) | chronic alcohol withdrawal severity |
|  | GS84278 | (Published QTL ) | chronic alcohol withdrawal severity |
|  | GS84279 | (Published QTL ) | METH responses for climbing |
| C1qa | GS86977 | [DRG] | Table S1: All transcripts significantly different in abundance between the majority of heroin subjects and their matched controls (provisional) |
|  | GS87058 | [DRG] | Table S2: Cocaine Regulation of H4 Acetylation. (provisional) |
|  | GS243385 | [MeSH] | Dose-Response Relationship, Drug : D004305 |
|  | GS235349 | [MeSH] | Physiological Effects of Drugs : D045505 |
|  | GS1243 | Differential Expression | KCgamma wild-type expression changes due to chronic ethanol diet |
|  | GS83985 | (Published QTL ) | cocaine related behavior 16 |
|  | GS14933 | Differential Expression | Upregulated gene expression of PKC-gamma wild type mice due to chronic ethanol diet |
|  | GS84164 | (Published QTL ) | cocaine related behavior |
|  | GS83998 | (Published QTL ) | cocaine and amphetamine-regulated transcript |
| Cp | GS87096 | [DRG] | Table S2: List of Cocaine-Treated HDAC5 KO vs. Cocaine-Treated WT Significantly Regulated Genes. (Provisional) |
|  | GS87041 | [DRG] | Table S3: List of Cocaine-Treated HDAC5 KO vs. Saline-Treated HDAC5 KO Significantly Regulated Genes. (provisional) |
|  | GS243385 | [MeSH] | Dose-Response Relationship, Drug : D004305 |
|  | GS235349 | [MeSH] | Physiological Effects of Drugs : D045505 |
| Btg2 | GS86789 | [DRG] | Table S1: Cocaine Regulation of H3 Acetylation. (provisional) |
|  | GS243385 | [MeSH] | Dose-Response Relationship, Drug : D004305 |
|  | GS235349 | [MeSH] | Physiological Effects of Drugs : D045505 |
|  | GS1243 | Differential Expression | KCgamma wild-type expression changes due to chronic ethanol diet |
|  | GS14933 | Differential Expression | Upregulated gene expression of PKC-gamma wild type mice due to chronic ethanol diet |
|  | GS37147 | Differential Expression | Gene expression change in the nucleus accumbens, following continuous alcohol consumption in alcohol preferring rats. |
| Cfb | GS87128 | [DRG] | Table S1: Genes with significant alterations in expression following acute nicotine treatment in VTA. |
|  | GS243385 | [MeSH] | Dose-Response Relationship, Drug : D004305 |
|  | GS235349 | [MeSH] | Physiological Effects of Drugs : D045505 |
|  | GS127417 | Differential Expression | Chronic alcohol exposure induced gene expression changes in the zebrafishbrain |
|  | GS84303 | (Published QTL ) | differences in cocaine responsiveness |
|  | GS84300 | (Published QTL ) | METH responses for body temperature |
|  | GS84303 | (Published QTL ) | differences in cocaine responsiveness |
|  | GS213106 | Differential Expression | Chronic Alcohol HepG2 |
|  | GS84301 | (Published QTL ) | ethanol conditioned taste aversion |
|  | GS83968 | (Published QTL ) | cocaine induced activation 13 |
|  | GS239299 | [MeSH] | Drug Interactions : D004347 |
|  | GS135650 | (Published QTL ) | cocaine induced activation 13 |
|  | GS84302 | (Published QTL ) | differences in cocaine responsiveness |
| Dnajb3 | GS87011 | [DRG] | Table S2: List of Cocaine-Treated HDAC5 KO vs. Cocaine-Treated WT Significantly Regulated Genes. |
|  | GS75590 | Differential Expression | Cocaine Regulation of Dimethyl-K9/K27 H3 |
|  | GS83978 | (Published QTL ) | cocaine related behavior 1 |
|  | GS83973 | (Published QTL ) | cocaine induced activation 5 |
|  | GS84103 | (Published QTL ) | chronic alcohol withdrawal severity Chr1 at D1Mit46 |
|  | GS135653 | (Published QTL ) | cocaine induced activation 5 |
|  | GS135293 | (Published QTL ) | alcohol withdrawal 5 |
| Hspb2 | GS87011 | [DRG] | Table S2: List of Cocaine-Treated HDAC5 KO vs. Cocaine-Treated WT Significantly Regulated Genes. |
|  | GS31782 | Differential Expression | Gene Expression Correlations with Hippocampus Consortium M430v2 (Jun06) RMA for Localization of genes affecting alcohol drinking in mice Phillips et al |
|  | GS235349 | [MeSH] | Physiological Effects of Drugs : D045505 |
| Il12rb2 | GS84180 | (Published QTL ) | METH responses for body temperature |
|  | GS84181 | (Published QTL ) | ethanol induced locomotion |
|  | GS84182 | (Published QTL ) | METH responses for home cage activity |
|  | GS84179 | (Published QTL ) | cocaine related behavior |
|  | GS135789 | (Published QTL ) | ethanol induced locomotor activity 2 |
|  | GS243385 | [MeSH] | Dose-Response Relationship, Drug : D004305 |
|  | GS83991 | (Published QTL ) | cocaine related behavior 7 |
|  | GS235349 | [MeSH] | Physiological Effects of Drugs : D045505 |
| Parp8 | GS128167 | Differential Expression | Table S1: Genes differentially expressed in Lewis vs. Fisher nucleus accumbens shell GABA projection neurons |
|  | GS246373 | Differential Expression | Differential Expression Hippocampus Human Alcoholic |
| Dnase1l2 | GS84300 | (Published QTL ) | METH responses for body temperature |
|  | GS84301 | (Published QTL ) | ethanol conditioned taste aversion |
|  | GS83968 | (Published QTL ) | cocaine induced activation 13 |
|  | GS83971 | (Published QTL ) | cocaine induced activation 3 |
|  | GS84298 | (Published QTL ) | cocaine induced activation |
|  | GS135650 | (Published QTL ) | cocaine induced activation 13 |
|  | GS84302 | (Published QTL ) | differences in cocaine responsiveness |
|  | GS36452 | Differential Expression | Whole Brain Gene expression correlates of Morphine - Postural Effects in Females & Males BXD |
|  | GS36477 | Differential Expression | Whole Brain Gene expression correlates of Morphine - Severity of ptosis inMales BXD |
|  | GS36457 | Differential Expression | Whole Brain Gene expression correlates of Morphine - Postural Effects in Females BXD |
| Htr1a | GS243385 | [MeSH] | Dose-Response Relationship, Drug : D004305 |
|  | GS235349 | [MeSH] | Physiological Effects of Drugs : D045505 |
|  | GS242550 | [MeSH] | Alcoholism : D000437 |
|  | GS324475 | GO | 0008144 drug binding |
|  | GS332669 | GO | 0017144 drug metabolic process |
|  | GS269429 | GWAS | Catalog Data for alcohol and nicotine codependence in 818 European ancestry cases, 1,396 European ancestry controls |
|  | GS246373 | Differential Expression | Differential Expression Hippocampus Human Alcoholic |
|  | GS236764 | [MeSH] | Drug-Related Side Effects and Adverse Reactions : D064420 |
|  | GS239299 | [MeSH] | Drug Interactions : D004347 |
|  | GS242397 | [MeSH] | Psychotropic Drugs : D011619 |
| Rilpl2 | GS135737 | (Published QTL ) | dopamine receptor binding 2 |
|  | GS136242 | (Published QTL ) | methamphetamine response QTL 1 |
|  | GS84174 | (Published QTL ) | METH responses for chewing |
|  | GS135490 | (Published QTL ) | behavioral response to methamphetamines 3 |
|  | GS135655 | (Published QTL ) | cocaine induced activation 7 |
|  | GS83975 | (Published QTL ) | cocaine induced activation 7 |
|  | GS84173 | (Published QTL ) | differences in cocaine responsiveness |
|  | GS84172 | (Published QTL ) | cocaine related behavior |
|  | GS246375 | Differential Expression | H3K4me3 CHiP Seq Hippocampus Human Alcoholics |
|  | GS84175 | (Published QTL ) | METH responses for climbing |
|  | GS84176 | (Published QTL ) | METH responses for climbing |
| Far2 | GS84189 | (Published QTL ) | ethanol conditioned taste aversion |
|  | GS127346 | Differential Expression | Transcripts differentially regulated in hippocampus of C57BL/6J mice drinking to intoxication. |
|  | GS84190 | (Published QTL ) | METH responses for body temperature |
| Pnmt | GS235349 | [MeSH] | Physiological Effects of Drugs : D045505 |
|  | GS136244 | (Published QTL ) | methamphetamine response QTL 3 |
|  | GS327758 | GO | 0035690 cellular response to drug |
|  | GS318209 | GO | 0017144 drug metabolic process |
|  | GS326077 | GO | 0042493 response to drug |
|  | GS135823 | (Published QTL ) | ethanol conditioned taste aversion 9 |
|  | GS37188 | (Published QTL ) | Positional candidate on Chromosome 11 (30-110 Mb) for dominant deviation measuring EtOH consumption during Drinking in the Dark (DID), 24 hour access and Blood Ethanol Concentration (BEC). |
| Cp | GS87096 | [DRG] | Table S2: List of Cocaine-Treated HDAC5 KO vs. Cocaine-Treated WT Significantly Regulated Genes. (Provisional) |
|  | GS87041 | [DRG] | Table S3: List of Cocaine-Treated HDAC5 KO vs. Saline-Treated HDAC5 KO Significantly Regulated Genes. (provisional) |
|  | GS243385 | [MeSH] | Dose-Response Relationship, Drug : D004305 |
|  | GS235349 | [MeSH] | Physiological Effects of Drugs : D045505 |
|  | GS84144 | (Published QTL ) | METH responses for home cage activity (Published QTL, Chr 3) |
|  | GS84146 | (Published QTL ) | METH responses for home cage activity (Published QTL, Chr 3) |
|  | GS128161 | Differential Expression | Nucleus accumbens Methamphetamine and reward |
|  | GS35864 | Differential Expression | Neocortex Gene expression correlates of Cocaine CPP - difference in percent test time spent relative to preconditioning in Females BXD |
|  | GS243385: | [MeSH] | Dose-Response Relationship, Drug : D004305 |
|  | GS86932 | [DRG] | Table S3: CORTEX 17K MICROARRAY |
|  | GS86494 | [DRG] | Table S3: CORTEX 17K MICROARRAY |
|  | GS243385 | [MeSH] | Dose-Response Relationship, Drug : D004305 |
|  | GS128199 |  | Alcohol Preference union of 86 Gene Sets |
|  | GS135133 | Differential Expression | bHR vs bLR genes different in Hippocampus |
|  | GS135132 | Differential Expression | bHR vs bLR genes different in Nucleus Acumbens |
| Dnaja4 | GS14888 | Differential Expression | Differentially expressed genes modulated by nicotine in five combined brain regions (Amygdala, Hippocampus, Nucleus Accumbens, Pre Frontal Cortex and Ventral Tegmental Area) for C3H/HeJ mice |
|  | GS135660 | (Published QTL ) | cocaine related behavior 8 (Cocrb8, Published QTL Chr 9) |
|  | GS135821 | (Published QTL ) | ethanol consumption 3 (Etohc3, Published QTL Chr 9) |
|  | GS135647 | (Published QTL ) | cocaine induced activation 10 (Cocia10, Published QTL Chr 9) |
|  | GS84219 | (Published QTL ) | cocaine related behavior (Published QTL, Chr 9) |
|  | GS84218 | (Published QTL ) | differences in cocaine responsiveness (Published QTL, Chr 9) |
|  | GS84217 | (Published QTL ) | differences in cocaine responsiveness (Published QTL, Chr 9) |
|  | GS84208 | (Published QTL ) | METH responses for home cage activity (Published QTL, Chr 9) |
|  | GS14914 | Differential Expression | Differentially expressed genes in morphine-treated vs. saline-treated peripheral blood mononuclear cells (PBMCs) |
|  | GS128199 |  | Alcohol Preference union of 86 Gene Sets |
|  | GS135133 | Differential Expression | bHR vs bLR genes different in Hippocampus |
|  | GS135132 | Differential Expression | bHR vs bLR genes different in Nucleus Acumbens |
| Dnmt3a | GS243385: | [MeSH] | Dose-Response Relationship, Drug : D004305 |
|  | GS86932 | [DRG] | Table S3: CORTEX 17K MICROARRAY |
|  | GS86494 | [DRG] | Table S3: CORTEX 17K MICROARRAY |
|  | GS243385 | [MeSH] | Dose-Response Relationship, Drug : D004305 |
|  | GS34054 | Differential Expression | Hippocampus Gene expression correlates of Open Field locomotion (cm) 45-60 min post cocaine in Males BXD |
|  | GS34322 | Differential Expression | Hippocampus Gene expression correlates of Cocaine TOTAL locomotion (activity beam breaks) in Males BXD |
|  | GS34005 | Differential Expression | Hippocampus Gene expression correlates of Open Field locomotion 15-30 min post cocaine in Males BXD |
|  | GS34332 | Differential Expression | Hippocampus Gene expression correlates of Cocaine TOTAL locomotion ( cm in 1 hr) in Males BXD |
|  | GS246394 | Differential Expression | Human hippocampus chronically exposed to cocaine |
|  | GS84261 | (Published QTL ) | ethanol withdrawal (Published QTL, Chr 12) |
|  | GS86746 | [DRG] | Table S5: List of Cocaine-Treated WT vs. Saline-Treated WT Significantly Regulated Genes. [DRG] |
|  | GS246374 | Differential Expression | Differential Expression Hippocampus Human Cocaine Addicts |
|  | GS128199 |  | Alcohol Preference union of 86 Gene Sets |
| Htr7 | GS243385: | [MeSH] | Dose-Response Relationship, Drug : D004305 |
|  | GS243385 | [MeSH] | Dose-Response Relationship, Drug : D004305 |
|  | GS242397 | [MeSH] | Psychotropic Drugs : D011619 |
|  | GS14914 | Differential Expression | Differentially expressed genes in morphine-treated vs. saline-treated peripheral blood mononuclear cells (PBMCs) |
|  | GS242550 | [MeSH] | Alcoholism : D000437 |
|  | GS84314 | (Published QTL ) | METH responses for body temperature (Published QTL, Chr 19) |
|  | GS236200 | [MeSH] | Neurotransmitter Uptake Inhibitors : D014179 |
|  | GS246374 | Differential Expression | Differential Expression Hippocampus Human Cocaine Addicts |
|  | GS128199 |  | Alcohol Preference union of 86 Gene Sets |
|  | GS135133 | Differential Expression | bHR vs bLR genes different in Hippocampus |
|  | GS135132 | Differential Expression | bHR vs bLR genes different in Nucleus Acumbens |
| Irf8 | GS243385: | [MeSH] | Dose-Response Relationship, Drug : D004305 |
|  | GS86932 | [DRG] | Table S3: CORTEX 17K MICROARRAY |
|  | GS86494 | [DRG] | Table S3: CORTEX 17K MICROARRAY |
|  | GS243385 | [MeSH] | Dose-Response Relationship, Drug : D004305 |
|  | GS1139 | Differential Expression | Differential expression response 4 hr after 2g/kg ethanol in C57BL/6J and DBA/2J |
|  | GS246374 | Differential Expression | Differential Expression Hippocampus Human Cocaine Addicts |
|  | GS128199 |  | Alcohol Preference union of 86 Gene Sets |
|  | GS135133 | Differential Expression | bHR vs bLR genes different in Hippocampus |
|  | GS135132 | Differential Expression | bHR vs bLR genes different in Nucleus Acumbens |
| Lrrc15 | GS84293 | (Published QTL ) | METH responses for home cage activity (Published QTL, Chr 16) |
|  | GS35781 | Differential Expression | Cerebellum Gene expression correlates of CPP - Time (s) in drug-paired compartment a in Males BXD |
| Myh10 | GS37188 | (Published QTL ) | Positional candidate on Chromosome 11 (30-110 Mb) for dominant deviation measuring EtOH consumption during Drinking in the Dark (DID), 24 hour access and Blood Ethanol Concentration (BEC). |
|  | GS84251 | (Published QTL ) | chronic alcohol withdrawal severity (Published QTL, Chr 11) |
|  | GS243385: | [MeSH] | Dose-Response Relationship, Drug : D004305 |
|  | GS135823 | (Published QTL ) | ethanol conditioned taste aversion 9 (Etohcta9, Published QTL Chr 11) |
|  | GS37187 | (Published QTL ) | Positional candidate on chromosome 11 (59-79Mb) for overdominant effect for 24-hour, 2 bottle choice 30g/kg EtOH excessive consumption. |
|  | GS86932 | [DRG] | Table S3: CORTEX 17K MICROARRAY |
|  | GS86494 | [DRG] | Table S3: CORTEX 17K MICROARRAY |
|  | GS243385 | [MeSH] | Dose-Response Relationship, Drug : D004305 |
|  | GS127342 | Differential Expression | Transcripts differentially regulated in frontal cortex of C57BL/6J mice drinking to intoxication. |
|  | GS75588 | Differential Expression | Cocaine Regulation of H3 Acetylation |
|  | GS137407 | Differential Expression | Supplementary Table 2. Overall results of WGCNA combined with differential expression between alcoholics and controls |
|  | GS313343 | Gene Ontology | GO:0008144 drug binding |
|  | GS36154 | Differential Expression | Neocortex Gene expression correlates of Locomotor response of 10 mg/kg MDMA injected on Day 2 in Females & Males BXD |
|  | GS75588 | Differential Expression | Cocaine Regulation of H3 Acetylation |
|  | GS128199 |  | Alcohol Preference union of 86 Gene Sets |
|  | GS135133 | Differential Expression | bHR vs bLR genes different in Hippocampus |
|  | GS135132 | Differential Expression | bHR vs bLR genes different in Nucleus Acumbens |
| Rap2b | GS84146 | (Published QTL ) | METH responses for home cage activity (Published QTL, Chr 3) |
|  | GS243385: | [MeSH] | Dose-Response Relationship, Drug : D004305 |
|  | GS84147 | (Published QTL ) | ethanol conditioned taste aversion (Published QTL, Chr 3) |
|  | GS86932 | [DRG] | Table S3: CORTEX 17K MICROARRAY |
|  | GS86494 | [DRG] | Table S3: CORTEX 17K MICROARRAY |
|  | GS243385 | [MeSH] | Dose-Response Relationship, Drug : D004305 |
|  | GS246373 | Differential Expression | Differential Expression Hippocampus Human Alcoholic |
|  | GS246394 | Differential Expression | Human hippocampus chronically exposed to cocaine |
|  | GS128199 |  | Alcohol Preference union of 86 Gene Sets |
| Tmod2 | GS35864 | Differential Expression | Neocortex Gene expression correlates of Cocaine CPP - difference in percent test time spent relative to preconditioning in Females BXD |
|  | GS14917 | Differential Expression | Upregulation of gene expression in the lateral hypothalamus of Wild Type (WT) mice following administration of chronic morphine |
|  | GS135660 | (Published QTL ) | cocaine related behavior 8 (Cocrb8, Published QTL Chr 9) |
|  | GS135821 | (Published QTL ) | ethanol consumption 3 (Etohc3, Published QTL Chr 9) |
|  | GS14916 | Differential Expression | Mu opioid receptor-dependent genes regulated by chronic morphine in the lateral hypothalamus (LH) |
|  | GS86932 | [DRG] | Table S3: CORTEX 17K MICROARRAY |
|  | GS86494 | [DRG] | Table S3: CORTEX 17K MICROARRAY |
|  | GS75567 | Differential Expression | Genes that were significantly different in the nucleus accumbens of iP rats between the ethanol and water groups |
|  | GS14929 | Differential Expression | Ethanol-dependence genes in the nucleus accumbens (NA) of inbred alcohol-preferring |
|  | GS75589 | Differential Expression | Cocaine Regulation of H4 Acetylation |
|  | GS137562 | Differential Expression | Genes significantly differentially expressed in P7 selected High-responder (bHR) vs. Low-responder (bLR) in the hippocampus of Sprague-Dawley rats. |
|  | GS128223 | Differential Expression | Proteins found to be modified by at least two drugs of abuse |
|  | GS135822 | (Published QTL ) | ethanol conditioned taste aversion 8 (Etohcta8, Published QTL Chr 9) |
|  | GS84219 | (Published QTL ) | cocaine related behavior (Published QTL, Chr 9) |
|  | GS84218 | (Published QTL ) | differences in cocaine responsiveness (Published QTL, Chr 9) |
|  | GS84217 | (Published QTL ) | differences in cocaine responsiveness (Published QTL, Chr 9) |
|  | GS246374 | Differential Expression | Differential Expression Hippocampus Human Cocaine Addicts |
|  | GS128199 |  | Alcohol Preference union of 86 Gene Sets |
|  | GS135133 | Differential Expression | bHR vs bLR genes different in Hippocampus |
|  | GS135132 | Differential Expression | bHR vs bLR genes different in Nucleus Acumbens |
| Zbtb4 | GS37188 | (Published QTL ) | Positional candidate on Chromosome 11 (30-110 Mb) for dominant deviation measuring EtOH consumption during Drinking in the Dark (DID), 24 hour access and Blood Ethanol Concentration (BEC). |
|  | GS84251 | (Published QTL ) | chronic alcohol withdrawal severity (Published QTL, Chr 11) |
|  | GS135823 | (Published QTL ) | ethanol conditioned taste aversion 9 (Etohcta9, Published QTL Chr 11) |
|  | GS37187 | (Published QTL ) | Positional candidate on chromosome 11 (59-79Mb) for overdominant effect for 24-hour, 2 bottle choice 30g/kg EtOH excessive consumption. |
|  | GS137413 | Differential Expression | Supplementary Table 2. CNA Overall results of WGCNA combined with differential expression between alcoholics and controls |
|  | GS246376 | Differential Expression | H3K4me3 CHiP Seq Hippocampus Human Cocaine Addicts |
|  | GS246376 | Differential Expression | H3K4me3 CHiP Seq Hippocampus Human Cocaine Addicts |
|  | GS246373 | Differential Expression | Differential Expression Hippocampus Human Alcoholic |
